# Multifaceted and evolutionarily dynamic interactions between *Caenorhabditis elegans* SPO-11 and its cofactors ensure proper formation of meiotic DNA double-strand breaks

**DOI:** 10.64898/2026.07.30.741518

**Authors:** Keita Kameda, Li Zhou, Nowa Matsubayashi, Carlos Mario Rodriguez-Reza, Lin Meng, Xuan Li, Tomoki Yoshioka, Aya Sato-Carlton, Peter Mark Carlton

## Abstract

DNA double-strand breaks (DSBs) generated during meiotic prophase by the topoisomerase-like protein SPO11 are essential to create crossovers between homologous chromosomes. Since crossovers are required to biorient chromosomes at the first meiotic division, DSB formation is essential for meiosis in most sexually-reproducing organisms. Since excess DSBs have the potential to destabilize the genome, SPO-11 activity must be strictly regulated by many cofactors. Recent studies have established that SPO11 must dimerize to cut DNA, whereas soluble SPO11 and SPO11-TOPOVIBL complexes are predominantly monomeric (1–3). This contrast suggested that a major role of SPO11 cofactors could be to promote SPO11 dimerization, through means such as increasing local concentration or co-orienting SPO11 protomers. However, the mechanism of this regulation is not well-understood. Here, by taking advantage of phylogenomic analysis in the nematode genus *Caenorhabditis*, we show that the conserved cofactor DSB-1^Rec114^ evolved to replace TOPOVIBL function in *C. elegans*. We provide genetic and biochemical evidence that multiple interactions between SPO-11 and DSB-1 stabilize protein complex formation and promote SPO-11 dimerization. Our results shed light on the regulatory mechanism of programmed DSB formation, which ensures crossover formation and meiotic chromosome segregation while protecting genomic stability.

**Significance Statement:** Programmed DNA double-strand breaks catalyzed by SPO11 are essential for meiosis, but how SPO11 and its cofactors cooperate to cut DNA is not understood. SPO11 only cuts DNA as a homodimer, but soluble SPO11, with or without its core component TOPOVIBL, is predominantly monomeric. We show here that DSB-1, a conserved cofactor of *C. elegans* SPO-11, has evolved to replace TOPOVIBL to make direct, multifaceted interactions with SPO-11. We provide evidence that DSB-1 simultaneously binds both SPO-11 protomers, and this binding is critical for DNA cleavage, implying a major role of DSB-1 in promoting SPO-11 dimerization. Our phylogenetic analysis also highlights the evolutionary flexibility of a conserved, essential protein complex after the loss of one of its members, TOPOVIBL.

## Introduction

Correct segregation of homologous chromosomes at meiosis requires crossover (CO) formation to establish biorientation of chromosomes at the first meiotic division. Failure in CO formation leads to chromosome nondisjunction, a major cause of aneuploid gametes leading to infertility (4–7). Crossover formation results from repair of programmed DNA double-strand breaks (DSBs) created by the topoisomerase-like protein Spo11, acting in concert with several cofactors (8–10). Spo11 is evolutionarily related to the A subunit of Topoisomerase VI (11, 12). The topoVI holoenzyme is composed of two TopoVIA subunits, possessing DNA cleavage activity, and two TopoVIB subunits, carrying the ATPase domain. Previous studies have identified TOPOVIBL (TopoVI subunit B-like protein), derived from the B subunit of topoVI, and have shown that a Spo11-TOPOVIBL complex is essential for meiotic DSB formation (13, 14). Similar to the DNA-cleaving domain in TopoVIA subunits, Spo11 breaks DNA by attacking a phosphodiester bond with a tyrosine residue OH, which becomes covalently attached to the newly-broken 5′ end. To generate the active site for the DNA cleavage reaction, two Spo11 molecules need to be placed in a symmetric manner so that each catalytic tyrosine can coordinate with residues and their complexed cations from their partner Spo11, to attack the phosphodiester backbone of each DNA strand (1, 2, 15, 16).

The initiation of DSBs must be strictly regulated, since an excess of damage could lead to apoptosis, mutation, or genome instability, while an insufficient number of DSBs could lead to a failure in crossover formation and thus non-disjunction of chromosomes at meiosis I (5, 6, 17–19). Therefore, SPO-11 activity must be regulated in space and time to achieve a sufficient but not excessive number of DSBs during meiotic prophase.

Recent biochemical studies have shown that although purified mouse SPO11 primarily exists as monomers in solution, SPO11-TOPOVIBL heterodimers or SPO11 alone at high concentration can dimerize and are capable of cleaving DNA *in vitro* (3). Analysis of the reaction kinetics in these studies showed that SPO11 dimerization is a key rate-limiting step regulating DNA cleavage activity, while most SPO11-TOPOVIBL proteins remain bound to DNA as an inactive monomer (or heterodimer of one SPO11 and one TOPOVIBL) in the *in vitro* reconstitution. In budding yeast, the function of TOPOVIBL in mouse is divided between Rec102, possessing distant homology to the TOPOVIBL transducer domain, and Rec104, which occupies the position of the GHKL domain (20, 21). Recent work has shown that Rec102 can make *cis-*contacts with its nearest Spo11 protomer, as well as *trans-*contacts with the protein Ski8 bound to the farther (across the DNA axis) Spo11 protomer, illustrating a potential mechanism of Spo11 dimerization (16).

Previous studies in mouse and *S. cerevisiae* on DSB cofactors have shown that the protein Rec114 acts in concert with Mei4 and Mer2/IHO1, together referred to as the RMM complex, to promote DSB initiation (22). Homologs of yeast Rec114 include mouse REC114 (23, 24) and *Caenorhabditis elegans* paralogs DSB-1 and DSB-2 (25), all of which are required for meiotic DSB formation. A homolog of Mei4, DSB-3, is also required for DSB formation in *C. elegans* (26), but no nematode homolog of Mer2/IHO1 has yet been identified.

The RMM complex has been shown to interact with SPO11-TOPOVIBL via direct binding of REC114 to TOPOVIBL (or Rec114 to Rec102/104 in budding yeast) (27, 28). Previous studies have also shown that the RMM complex can form liquid-like condensates *in vitro*, which together with the helical periodicity observed in concerted Spo11 "double cuts", suggests that RMM condensates can act as a scaffold to concentrate and co-orient Spo11 proteins *in vivo* (29).

While SPO11 is highly conserved in sexually-reproducing eukaryotes, TOPOVIBL shows remarkable evolutionary divergence across species (30). As noted for yeast Rec102 above, many species have lost portions of TOPOVIBL, or otherwise retain only highly truncated versions of the gene. While ancestral TOPOVI requires ATP-driven conformational changes for religation of the cleaved DNA ends, programmed DSB formation during meiosis requires that DNA ends remain cleaved to serve as substrates for repair to initiate crossover formation. This biological difference may have led to diversity and degeneration in the ATP-binding domain of TOPOVIBL during evolution. A previous study (30) has shown that many nematodes, including *C. elegans,* entirely lack a recognizable TOPOVIBL. Since TOPOVIBL is required for DSBs, and the C-terminus of TOPOVIBL mediates interactions between SPO11 and the RMM complex in other model organisms (27), the absence of TOPOVIBL in some nematodes raises the question of what adaptations have allowed meiotic DSB formation without TOPOVIBL.

Previous yeast two hybrid (Y2H) assays suggested that SPO-11 can directly bind to DSB-1^Rec114^ but not to DSB-2^Rec114^ (31) whereas DSB-1^Rec114^, DSB-2^Rec114^ and DSB-3^Mei4^ are predicted to form a stable heterotrimer similar to the Rec114-Rec114-Mei4 trimer observed in other organisms (32–34). However, evidence for the overall structure of the SPO-11 and RMM complex components (DSB-1, 2, 3) in the DSB machinery is still lacking in *C. elegans*.

Our previous work has shown that duplication of an ancestral Rec114 gene into the paralogs DSB-1 and DSB-2 happened early in the *Caenorhabditis* genus (35). Known Rec114 homologs, including both DSB-1 and DSB-2, contain a conserved N-terminal PH domain, an alpha-helical C-terminus that engages in trimer formation, and an intrinsically disordered region (IDR) in between. DSB-1 is absolutely required for DSBs, and *dsb-1* null mutants mimic *spo-11* loss-of-function mutants (36). In contrast, loss of *dsb-2* still allows a reduced level of DSB initiation, which declines further with maternal age (37). Since DSBs are not completely eliminated in *dsb-2* mutants, DSB-1 on its own can suffice to promote the break activity of SPO-11, although this activity is lower compared to when DSB-2 is present. Previously, we have shown that DSB-1 activity is regulated by phosphorylation and dephosphorylation in an ATR kinase and PP4 phosphatase-dependent manner, and proposed that unphosphorylated DSB-1 promotes SPO-11’s DNA cleavage activity while phosphorylated DSB-1 inhibits DNA cleavage. In contrast to this regulation of DSB-1, DSB-2 appears insensitive to phosphoregulation by ATR kinase, highlighting another difference between these paralogs (35). Since the mode of DSB-1, DSB-2, and DSB-3 interaction with SPO-11 is not understood well, the mechanism by which this complex regulates SPO-11’s activity remains an unsolved mystery.

*C. elegans* as an experimental model has provided many new insights into meiosis. As of this writing, the genomes of more than 50 *Caenorhabditis* nematodes have been sequenced, annotated, and made freely available, creating an excellent resource for comparative genomics and evolutionary biology. This wide breadth of genomic information in combination with phylogenetic analysis established in nematodes lets us compare and identify correlation or coevolution of genes and functional protein domains during the evolution of nematode lineages.

In this study, in order to find connections between the putative DSB-1-2-3 complex and SPO-11, we searched for conserved protein domains correlating with loss of *top6bl* and found a conserved motif in the DSB-1^REC114^ PH domain, which is predicted by AlphaFold3 (AF) (38) to bind directly to the SPO-11 N-terminal winged helix domain. Interestingly, AF also predicted that one DSB-1 molecule can simultaneously bind to two SPO-11 molecules at multiple surfaces. We show that these predicted interaction sites can bind to each other *in vitro*, and mutations to the predicted interacting surface between DSB-1 and SPO-11 lead to reduction in DSB formation. In addition, our biochemical work also has shown that the SPO-11 N-terminus as well as a *Caenorhabditis*-specific C-terminal extension region in SPO-11 can bind to the PH domains of both DSB-1 and DSB-2 *in vitro*. By taking advantage of the availability of genomic sequence data in nematodes and comparative sequence alignments, our work suggests that DSB-1 evolved to replace TOPOVIBL function, and that multifaceted interactions between SPO-11 and DSB-1-2-3 promote SPO-11 dimerization and thus break formation without TOPOVIBL. Our results shed light on the regulatory mechanism of programmed DSB formation, which ensures crossover formation and meiotic chromosome segregation while protecting genomic stability.

## Results

### Emergence and conservation of motifs in DSB-1 correlating with loss of *top6bl* in the nematode lineage

The gene *top6bl* has evolved rapidly, losing various functional domains in different clades (30). A previous phylogenetic analysis showed that: (1) ancestral nematodes possess a *top6bl* gene, but (2) in 13 out of 20 diverse nematode species, including *C. elegans*, *top6bl* is undetectable. The absence of *top6bl* was suggested to correlate with an nematode-specific extension of the *spo-11 C*-terminus (30). The coincidence of losing *top6bl* and acquiring the *spo-11* C-terminus extension during *Caenorhabditis* evolution raised the possibility that the SPO-11 C-terminus extension can functionally replace TOPOVIBL or enable SPO-11 to interact with a novel mediator bridging SPO-11 and the RMM complex.

However, when we analyzed multiple sequence alignments (MSAs) of SPO-11, DSB-1 and DSB-2 sequences of 54 species of *Caenorhabditis* for more extensive comparison, we found that a set of 17 closely-related species diverged after loss of the *top6bl* gene but before emergence of the SPO-11 C-terminus extension (**Figure 1A, 1B**). This shows that the SPO-11 C-terminus extension is not strictly anticorrelated with *top6bl*. We therefore searched for other protein motifs correlating with loss of *top6bl* among DSB-related genes, and found that the emergence of the conserved amino acid motif [F/Y]ISE[S/T]PXK in the DSB-1^REC114^ PH domain correlated extremely well with loss of *top6bl* (**Figure 1C**). We found that 50 out of 54 *Caenorhabditis* species, including *C. elegans*, lack *top6bl* and possess the conserved *dsb-1* [F/Y]ISE[S/T]PXK motif, whereas the three most basal species (*C. parvicauda, C. auriculariae* and *C. monodelphis)* that have retained *top6bl* do not possess this motif in their *dsb-1^rec114^*genes (**Figure 1C**).

**Figure 1:**
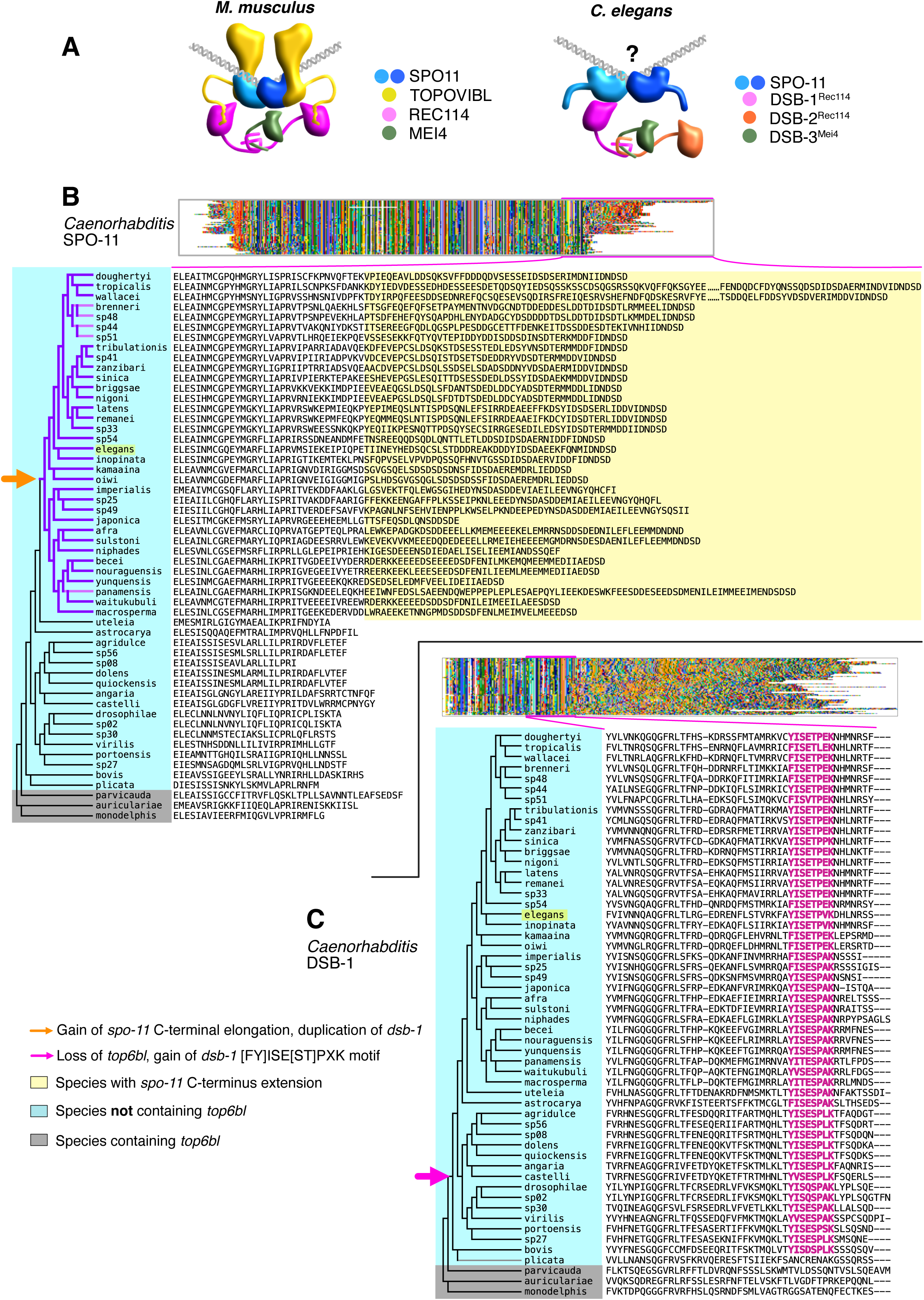
Emergence of the DSB-1 [FY]ISE[ST]PXK motifs correlates with loss of *top6bl* in the nematode lineage **(A)** Schematic diagrams of DSB initiation complex components in *M. musculus* (left) and *C. elegans* (right). **(B)** (Top) Multiple sequence alignment (MSA) image of SPO-11 proteins (entire length, 1 pixel per residue) in the order of the species phylogenetic tree below. (Bottom left) Species phylogenetic tree of 54 *Caenorhabditis* species (left), including *C. elegans* (yellow highlight), and MSA of their SPO-11 proteins at the C-terminus (right) of the corresponding worm. Worms with the *top6bl* gene are colored in grey, and worms lacking *top6bl* are colored in blue. The extended C-terminal region is colored in yellow. The phylogenetic branch at which the extended C-terminal region emerged is indicated by the orange triangle and darker branches. Darker purple lines in the phylogenetic tree indicate the species with DSB-2, and light pink lines indicate the ones which have lost DSB-2. **(C)** (Top) MSA image of DSB-1 proteins (entire length) in the order of the species phylogenetic tree below. (Bottom left) Species phylogenetic tree of 54 *Caenorhabditis* species (left), including *C. elegans* (yellow highlight), and MSA of their DSB-1 proteins at the boxed region (right). The conserved [FY]ISE[ST]PXK motifs are denoted in magenta, and the phylogenetic branch at which *top6bl* was lost and the [FY]ISE[ST]PXK motifs were gained is shown by the arrow in magenta. Conservation of this motif in DSB-1 correlates with loss of TOPOVIBL.

Examination of the gene distribution on the phylogenetic tree shows that *top6bl* was most likely lost just after the divergence of *C. parvicauda*; the early-diverging sister species *C. bovis* and *C. plicata* also lack *top6bl*, and *C. bovis* possesses the DSB-1 motif. The strong correlation between loss of *top6bl* and emergence of the conserved motif in *dsb-1* raised the possibility that the DSB-1 [F/Y]ISET[S/T]PXK motif has taken over the role of ancestral TOPOVIBL, promoting SPO-11 dimerization.

Since loss of *top6bl* did not correlate with the *spo-11* C-terminal extension, we next searched for other sequence features correlating with the emergence of the C-terminal extension of *spo-11* among DSB-related genes using multiple sequence alignment (MSA). We found that acquisition of an extended *spo-11* C-terminus co-occurred with duplication of the ancestral *dsb-1* gene (**Supplemental Figure S1, S2**) that gave rise to DSB-2. As we previously reported, the duplication of ancestral *dsb-1* happened when the lineage leading to the Elegans and Japonica groups (monophyletic subgroups of genus *Caenorhabditis*) diverged (35). All Elegans and Japonica species surveyed retain a *spo-11 C*-terminus extension, and no species outside these groups possesses it. This strong correlation raises the possibility of a functional relationship between the extended SPO-11 C-terminus and DSB-2. In both the Elegans and Japonica groups, the DSB-2 paralog has lost the [F/Y]ISE[S/T]PXK motif (**Supplemental Figure S2A**). Taken together, our phylogenetic analysis shows two notable coevolutionary changes in the *Caenorhabditis* DSB initiation machinery: first, the loss of *top6bl* with concomitant gain of the DSB-1 [F/Y]ISE[S/T]PXK motif, and second, the extension of the *spo-11* C-terminus with concomitant duplication of ancestral *dsb-1*.

### Direct and multifaceted interactions between DSB-1^REC114^ and SPO-11 are predicted

Since our phylogenetic analysis suggested a possible relationship between loss of *top6bl* and the [FY]ISE[ST]PXK motif (YISETPVK in *C. elegans*) in *dsb-1*, we first examined whether *C. elegans* DSB-1 can bind to SPO-11 directly via its YISETPVK motif. AlphaFold3 (AF) consistently shows *C. elegans* DSB-1/2/3, SPO-11 and DNA with four Mg^2+^ ions are predicted to form a complex, poising two SPO-11 molecules in an orientation complexed with DNA that is underwound and bent at 124.2°, with the OH of catalytic tyrosine residues usually within 4Å of phosphodiester bonds whose transesterification would result in the expected 2-base 5′ overhang products, similar to predicted and observed SPO11-DNA complexes from other model organisms (2, 15, 16) (**Figure 2A**). To analyze AF predictions quantitatively, we developed a tool to show predicted protein-protein interactions and their frequencies as a circos plot (see Materials and Methods) based on repeated AF modelling. Predicted interactions with high frequency (more than ten times out of twenty predictions) are shown in **Figure 2B**. In this model, a direct interaction between the YISETPVK (102-109aa) motif of DSB-1 and the nearest (*cis-*)SPO-11 molecule (49-57aa) is predicted with high frequency (**Figure 2B**: YISETPVK motif is underlined). To assess the confidence of this prediction, we generated further AF predictions of the winged-helix domain of SPO-11 (37-154aa) with the PH domain of DSB-1 (6-130aa) in isolation, which strongly support binding between them (iPTM=0.91, PTM=0.93, mean pLDDT=92.2). In contrast, binding of the PH domain of DSB-2 (7-131aa) to the SPO-11 winged helix domain in isolation was not predicted to occur (iPTM=0.18, PTM=0.53, mean pLDDT=75.0). In the full complex, DSB-1 is also predicted to contact the same *cis-*SPO-11 at two other less-conserved regions: DSB-1 residues 59-66 (VSRYPSLK) and 121-125 (TDVWY). Interestingly, AF predicted that DSB-1 residue R116 also interacts with the farther (*trans-*)SPO-11 protomer in the full complex (**Figure 2B**: magenta line in the circos plot), suggesting a single DSB-1 may interact with both poised SPO-11 protomers at the same time. Repeated AF modelling consistently predicted the conserved YISETPVK (102-109aa) sequence in DSB-1 binds to the *cis*-SPO-11 in nearly 100% of models, while DSB-1 R116 residue flips out to bind to the *trans-*SPO-11 molecule in roughly 50% of models. This prediction suggested the hypothesis that multifaceted interactions between DSB-1 and two SPO-11 molecules may stabilize the formation of active SPO-11 dimers. The predicted DSB-1 interaction surface on SPO-11, RKIEFALAD (49-57aa), corresponds spatially to the region where mouse SPO11 is predicted to bind to mouse TOPOVIBL by AF, suggesting that DSB-1 evolved to replace the ancestral TOPOVIBL binding at the same SPO-11 surface (**Figure 2C**). This SPO-11 motif (RKIEFALAD) is conserved across a wide range of nematodes, including the majority of nematodes that have retained *top6bl*, and is notably similar to the mammalian sequence (30).

**Figure 2:**
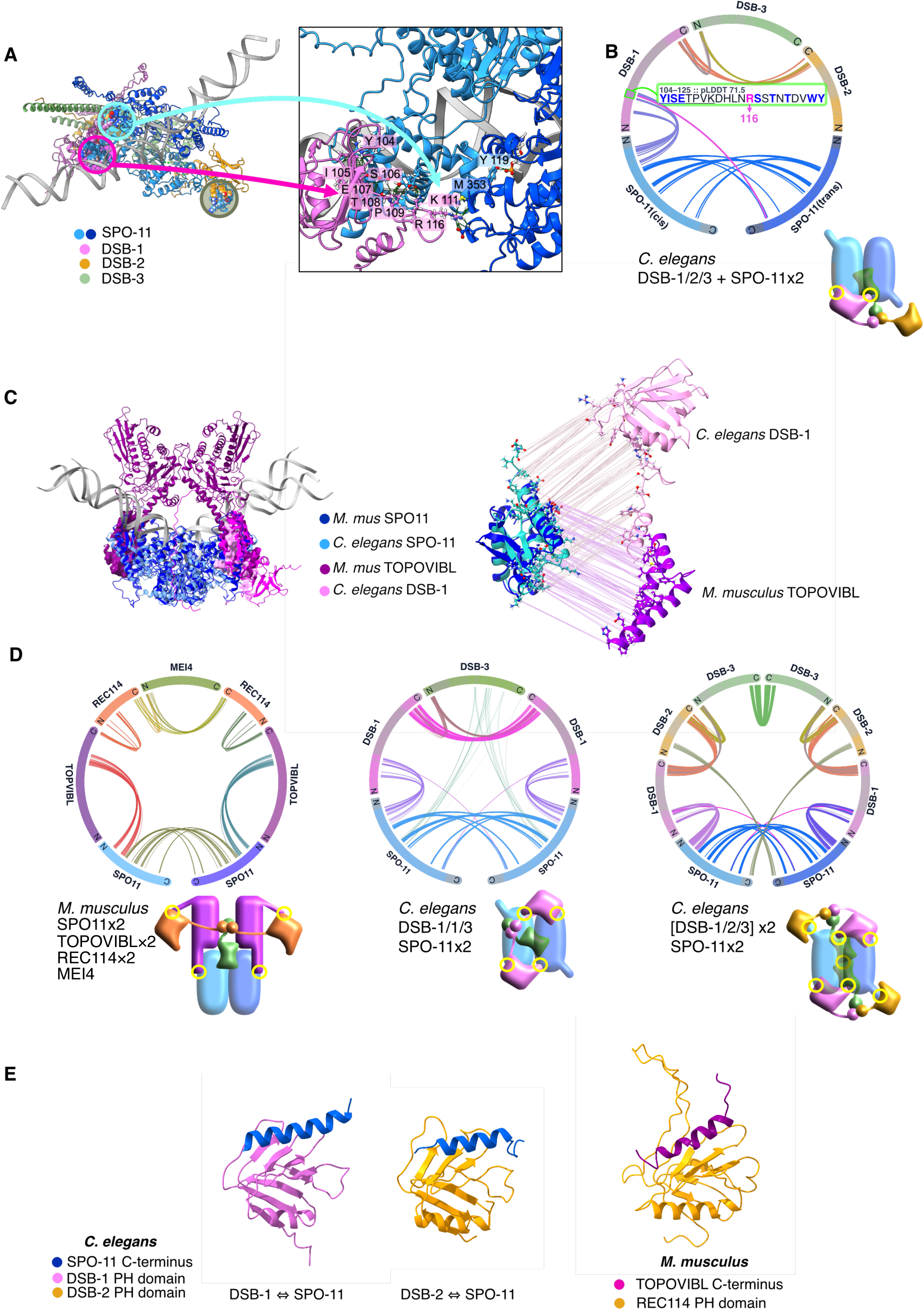
Direct and multifaceted interactions between DSB-1^REC114^ and SPO-11 are predicted. (A) AlphaFold3 (AF) prediction of protein complex of *C. elegans* SPO-11×2, DSB-1/2/3 and DNA (left), and its enlarged image (right) showing the DSB-1 [FY]ISE[ST]PXK motif (magenta circle) contacting *cis*-SPO-11, DSB-1 R116 (cyan circle) contacting *trans-*SPO-11, and DSB-2 interacting with a SPO-11 C terminus (brown circle). DNA with a single-strand nick is used for prediction, which serves to orient the protein complexes consistently in each model. (B) Circos plot of contacts calculated by ChimeraX on AF3-predicted protein complexes of *C. elegans* SPO-11×2, DSB-1/2/3 and DNA (DNA is not shown). For each protein in the plot, angular ends denote the N-terminus while round ends denote the C-terminus of the protein. AF predictions are repeated four times (total 20 predictions), and the predicted interactions detected by ChimeraX more than half of the time (10 times or more out of 20 predictions) are shown. The amino acid sequence (104-125aa) in DSB-1 is shown in the green box, with the YISETPVK motif underlined in black, R116 residue indicated in magenta, and residues predicted to directly contact with SPO-11 shown in blue. The interaction between DSB-1 R116 and *trans*-SPO-11 is shown by a magenta line. The cartoon shows a modeled interpretation of the circos plot, with yellow circles indicating predicted sites of contact. (C) (Left) Overlay of AF predicted structures of the *M. musculus* SPO11×2 and TOPOVIBL complex and the *C. elegans* SPO-11×2 and DSB-1 complex. Ribbon models omit predicted IDR regions of SPO-11 and DSB-1; surfaces shown are TOPOVIBL and DSB-1 predicted contact surfaces. (Right) Predicted atomic interactions drawn as lines between *M. musculus* SPO11 and TOPOVIBL as well as *C. elegans* SPO-11 and DSB-1 in the indicated colors, showing the SPO-11 surface is common for both interactions. (D) Circos plots of AF-predicted protein interactions with the threshold of 10 or more positive interactions out of 20 predictions is shown. All predictions include the same nicked dsDNA (see Methods). (Left) *M. mus* complex of SPO11×2, TOPOVIBL, REC114×2, and MEI4, (Middle) *C. elegans* SPO-11×2, DSB-1×2 and DSB-3, (Right) *C. elegans* SPO-11×2, [DSB-1, DSB-2, DSB-3]×2 (octamer). Cartoons show modeled interpretations of circos plots, with yellow circles indicating predicted sites of contact. (E) AF-predicted protein interactions between PH domains of *C. elegans* DSB-1, DSB-2 with the SPO-11 C-terminal extension region (Left, Middle) and between the PH domain of *M. musculus* REC114 and the TOPVIBL C-terminus (558–579aa) (Right). All PH domains were first aligned to each other via ChimeraX "matchmaker" command, then the three models were separated by translation in the X axis, maintaining theirorientations.

In the AF models of the mouse TOPOVIBL×2, REC114×2, MEI4 and SPO11×2 complex with DNA, two TOPOVIBL molecules individually bind to two SPO11 molecules as well as two REC114s, while the REC114 and MEI4 subunits form a 2:1 trimer at the REC114 C-terminus and MEI4 N-terminus, creating a symmetric protein complex (**Figure 2D** **left**). In contrast, AF modeling of *C. elegans* DSB-1-2-3, SPO-11×2 and DNA predicted interactions between DSB-1 and SPO-11 with high frequency and likelihood (20/20 models, contact probabilities 0.66 ∼ 0.96) but interactions between DSB-2 and SPO-11 were predicted with low frequency and in different places, leading to an asymmetric protein complex prediction (**Figure 2B**).

Since DSB formation occurs, albeit at reduced levels, in *C. elegans dsb-2* mutants, and is expected to occur normally in other *Caenorhabditis* nematodes that naturally lack *dsb-2*, a protein complex of DSB-1-1-3 and SPO-11×2 should be able to generate DSBs in the absence of DSB-2. AF predicts that a protein complex of DSB-1-1-3 (DSB-1×2, DSB-3), SPO-11×2 and DNA can also be formed (**Figure 2D** **middle**), leading to a symmetric complex similar to the one in mice or yeast, but the trimerization of DSB-1-1-3 is predicted with lower frequency, suggesting that the DSB-1-1-3 trimerization is less stable than that of DSB-1-2-3. AF did not predict complex formation of [DSB-2-2-3, SPO-11×2, DNA] nor trimerization of [(DSB-2)×2 and DSB-3] by itself, consistent with the previous observation that DSB-1 is essential for DSB formation while DSB-2 is not absolutely essential (36, 37).

Considering that a putative asymmetric *C. elegans* complex of [DSB-1-2-3, SPO-11×2 and DNA] would leave empty one potential SPO-11/DSB-1 binding valency, we explored different protein stoichiometries and found that AF can also predict higher-order [(DSB-1-2-3)×2, SPO-11×2] complexes with DNA (**Figure 2D** **right**). This octamer complex restores the symmetry of the protein complex similar to the mouse model of [REC114×2, MEI4, SPO-11×2] with DNA, and raises the possibility that there may be multiple forms of DSB machinery complex containing DSB-1/2/3 and SPO-11 in *C. elegans*. Interestingly, in this octamer model of [(DSB-1-2-3)×2, SPO-11×2] with DNA, both DSB-1 molecules bind to both SPO-11 molecules, and DSB-2 is predicted to bind to the extended C-terminus of SPO-11 (**Figure 2D** **right**), while DSB-3 is strongly predicted to dimerize via its C-terminus.

Since functional relevance between DSB-2 and the SPO-11 C-terminus extended region (405-425aa) was suggested from the phylogenetic analysis as above, we also specifically examined if the SPO-11 C-terminus interacts with DSB-2 or DSB-1 in the predicted pentamer or octamer complexes [DSB-1-2-3, SPO-11×2]. The SPO-11 extended C-terminus contains many acidic residues towards the end of the protein from residues 405-425, and AF predicted its interaction with mostly basic residues in the PH domains of DSB-1 (residues 16–24 and 62–84) and DSB-2 (residues 12–17 and 53–80). This predicted binding surface on DSB-1 is different from the aforementioned, conserved motif YISETPEK (102-109aa), but is similar to binding between the C-terminus of mouse TOPOVIBL and REC114 (**Figure. 2E**). Taken together, at least three direct interactions are predicted by AF between DSB-1/2 and SPO-11, including (1) DSB-1 YISETPVK (102-109aa) motif with *cis*-SPO-11, (2) DSB-1 R116 residue with *trans-*SPO-11, and (3) SPO-11 extended C-terminus with DSB-1 or DSB-2.

### Direct interaction of DSB-1^REC114^ PH domain and SPO-11 N-terminus and C-terminal extension *in vitro*

Based on the AF predictions, we assessed possible protein-protein interactions between SPO-11 and DSB-1/2 with a pulldown assay. We designed a maltose-binding protein (MBP)-tagged SPO-11 N-terminus (31-150aa) construct to contain the RKIEFALAD (49-57aa) motif, predicted to bind to the DSB-1 YISETPVK motif, and a MBP-SPO-11 C-terminus construct containing the C-terminal acidic patch (405-425 aa)(**Figure 3A**). Both MBP-tagged SPO-11 constructs were able to pull down a 10xHis-tagged DSB-1 PH domain (1-112aa) containing the conserved YISETPVK motif as well as a 10xHis-tagged DSB-2 PH domain (1-108 aa) whereas SPO-11 did not interact with the negative control protein 10xHis-tagged HTP-3 (409-739 aa) (**Figure 3B**, **Supplemental Figure S3A**). Although AF predicts structures with the SPO-11 N-terminus bound to DSB-1 much more often than to DSB-2, our pull down assays showed that the SPO-11 N-terminus can bind to the DSB-2 PH domain.

**Figure 3:**
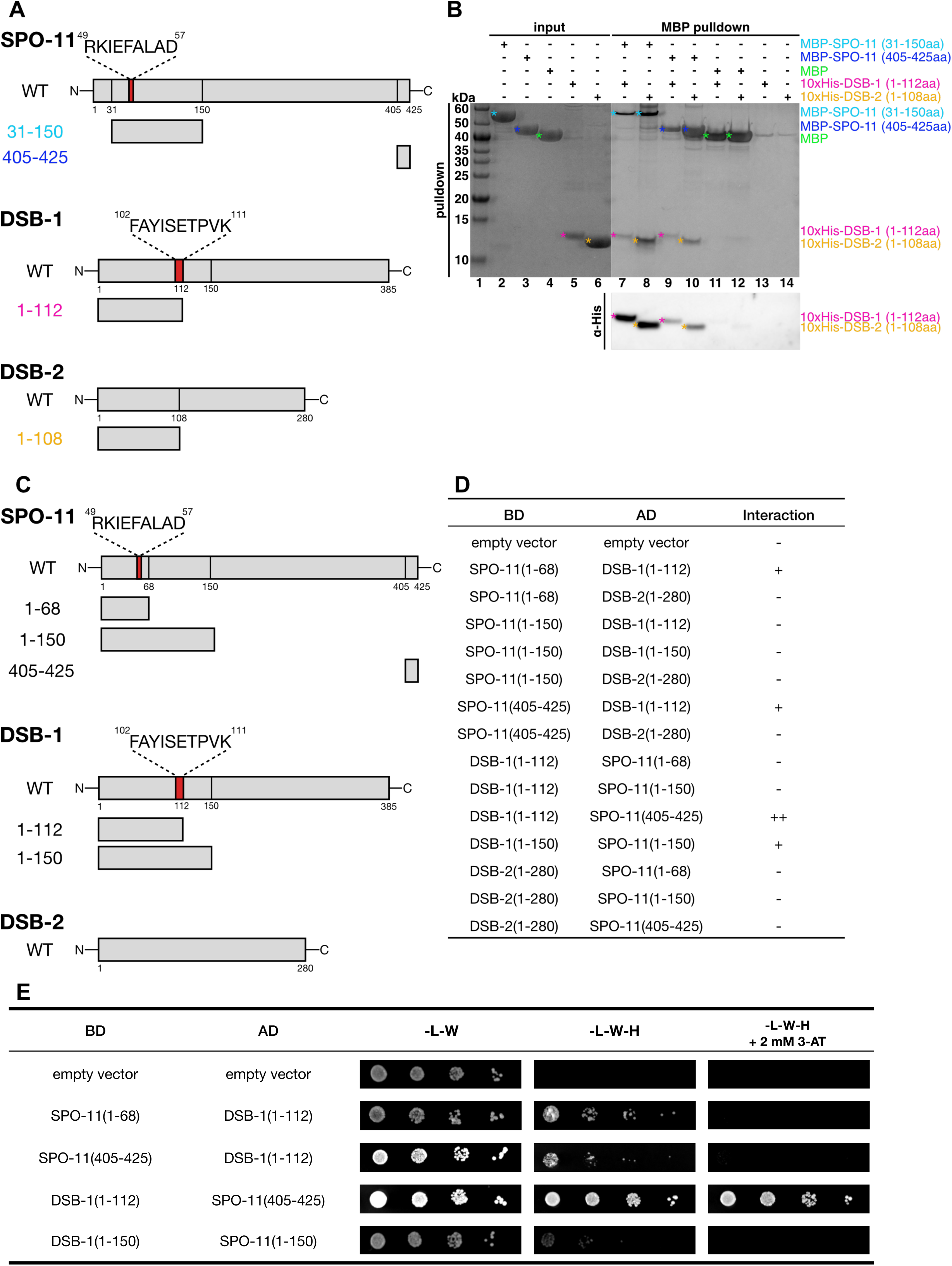
DSB-1^REC114^ PH domain directly interacts with the N terminus and C-terminal extension of SPO-11 *in vitro* **(A)** Schematic representation of the protein fragments used for the pulldown assay. SPO-11 fragments (31-150aa or 405-425aa) were fused to MBP-tag, whereas DSB-1 (1-112aa) and DSB-2 (1-108aa) fragments were fused to 10xHis-tag. **(B)** (Top)10xHis-tagged DSB-1 PH domain (1-112aa) and 10xHis-tagged DSB-2 PH domain (1-108aa) were pulled down by the MBP-tagged SPO-11 N-terminal (lane 7 and 9) or C-terminal (lane 8 and 10) fragments but not by the MBP control. Proteins in the input (lanes 1-6) were visualized by Coomassie Brilliant Blue staining and proteins in the eluates (lanes 7-14) were visualized by silver staining. (Bottom) Pulldown elutions were used for Western blot using α-His-tag antibodies to detect 10xHis-DSB-1 or DSB-2. **(C)** Schematic representation of the protein fragments used for the yeast two-hybrid (Y2H) assays. Proteins were fused to either the GAL4 DNA binding domain (BD) or the activation domain (AD). **(D)** Summary of the representative Y2H results from 10-fold to 10^4^-fold serial dilutions. Growth was assessed on medium lacking leucine (L) and tryptophan(W) and/or histidine (H), and with 2 mM 3-amino-1,2,4-trizole (3-AT). Since full-length DSB-1 fused to either BD or AD as well as BD-DSB-2 PH domain (1-108) exhibited self-activation, these constructs were excluded from the assay. **(E)** Representative Y2H results with serial dilution of cell number inoculation (10 to 10^7^-fold dilution) for the positive interactions summarised in **(D)** are shown. The N-or C-terminal fragment of SPO-11 directly bound to the DSB-1 PH domain.

We further examined the protein-protein interaction by Y2H assays. Two previous studies using Y2H assays have shown that SPO-11 full length binds to DSB-1 full length (31) and a SPO-11 fragment (48-425aa) binds to full length DSB-1 (26). In the latter study, their SPO-11 construct (Δ1-47aa) was clipped off near the RKIEFALAD (49-57aa) motif, predicted to interact with DSB-1, due to self-activation. In order to assess the SPO-11/DSB-1 or DSB-2 binding at the predicted domains, we prepared two SPO-11 N-terminal fragments: SPO-11 (1-68aa) and (1-150aa), one SPO-11 C-terminal fragment: SPO-11 (405-425aa), two DSB-1 N-terminal fragments : core PH domain (1-112aa), and the N-terminus fragment (1-150aa) containing the core PH domain as well as R116, and one DSB-2 full length construct (**Figure 3C**). The yeast constructs carrying either DSB-1 full length (1-385 aa) or DSB-2 PH domain only (1-108 aa) led to self-activation in our hands and thus were removed from the analysis. Positive Y2H interactions are summarized in **Figure 3D** and **3E**.

These Y2H data further support the hypothesis that SPO-11 N-terminus and C-terminus can directly interact with the DSB-1 PH domain. Many of the tested combinations, including the ones shown to be positively interacting in pull down assays, were negative in Y2H assays (**Supplemental Figure S3B**) possibly because protein folding of cloned constructs was not correct in yeast cells.

### Mutations in *spo-11* at the predicted DSB-1-SPO-11 interaction surface lead to reduction in DSB formation *in vivo*

To examine the functional importance of these possible interaction surfaces between DSB-1 and SPO-11 *in vivo*, we either introduced point mutations to three conserved residues often predicted to interact with conserved DSB-1 residues at the SPO-11 N-terminus (SPO-11 R49A, E52A, F53A: hereafter 3A mutant), or deleted the C-terminus extension region of SPO-11 to disrupt binding to DSB-1 and/or DSB-2. Since AF predicted that the terminal 21 residues (SPO-11 405-425aa) within the C-terminal extended region (SPO-11 371-425aa) binds to DSB-1/2 PH domains, we generated two different *spo-11* deletion mutants: the *spo-11(ΔC21)* mutant, deleting only the C-terminal acidic patch, and the *spo-11(ΔC39)* mutant, deleting the entire C-terminal extension region in the N2 wild type background (**Figure 4A**).

**Figure 4:**
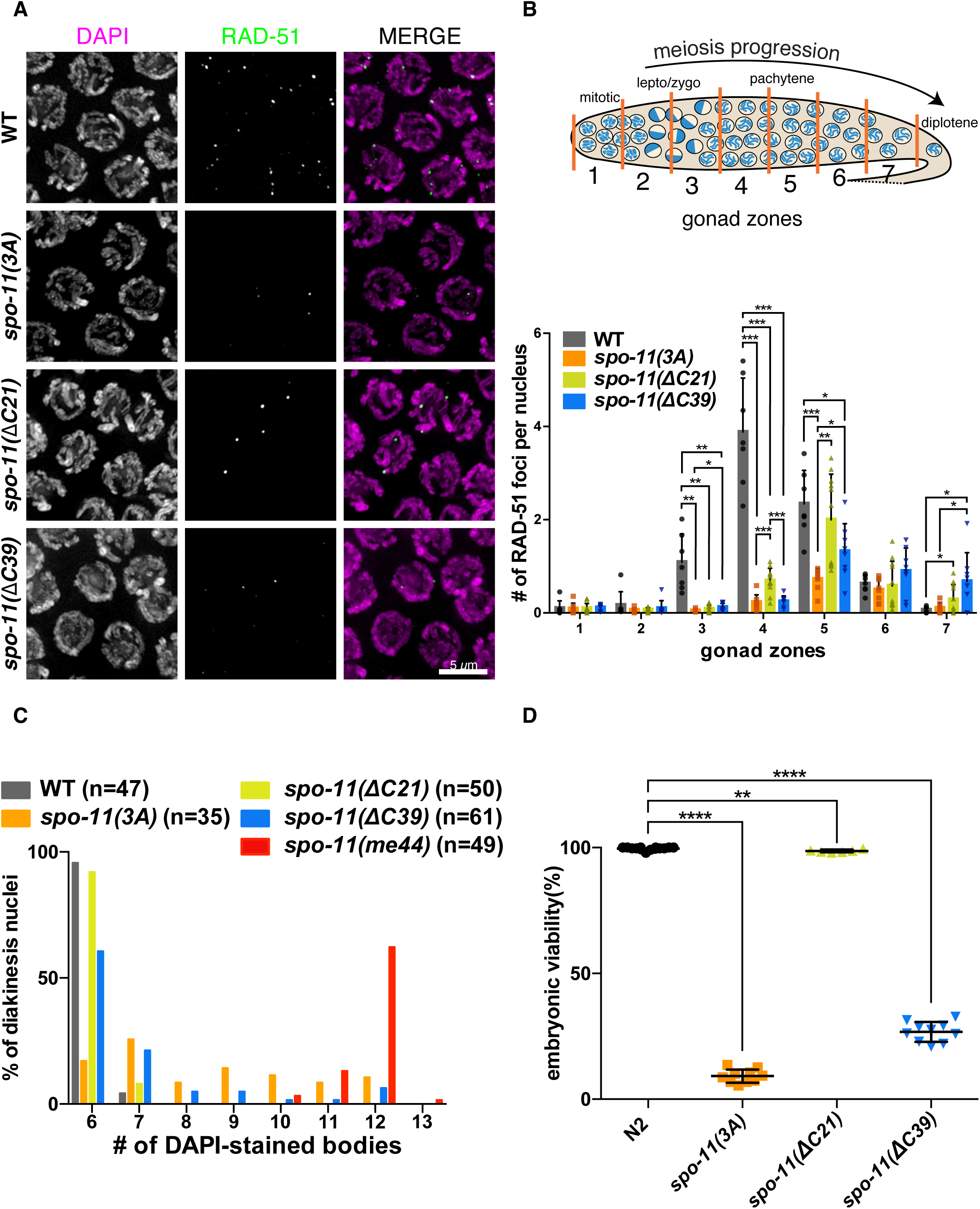
SPO-11 mutations at the predicted DSB-1 interaction surface reduced DSB production in oocytes (A) Representative immunofluorescence images of RAD-51 foci in mid-pachytene oocytes (zone 4) of the gonads of control (N2), *spo-11(3A), spo-11(ΔC21)* and *spo-11(ΔC39)* mutants. Scale bar, 5 µm. **(B)** (Top) Schematic representation of a hermaphrodite gonad divided into seven equal-length zones for RAD-51 focus scoring. (Bottom) Quantification of RAD-51 foci in the gonads of indicated genotypes in **(A)**. Data are presented as mean±SD (bar, overall mean; dots, mean of each individual gonad); The numbers of scored nuclei in zone 1-7 were as follows: for wild type (7 gonads), 348, 472, 488, 570, 468, 337, 120; for *spo-11(3A)* (8 gonads), 398, 681, 743, 726, 619, 404, 110; for *spo-11(ΔC21)* (11 gonads), 646, 850, 783, 891, 732, 521, 140; for *spo-11(ΔC39)* (8 gonads), 397, 586, 589, 610, 553, 387, 120. Statistical significance was assessed using a two-tailed t-test with Welch’s correction, **p<0.01, ***p<0.001. **(C)** Distribution of DAPI-stained bodies in diakinesis oocytes, shown as percentages, for the indicated genotypes. **(D)** Embryonic viability of the self-fertilized progeny of hermaphrodites with indicated genotypes. Data are presented as percentages. Statistical significance was assessed using a two-tailed t-test with Welch’s correction, **p<0.01, ****p<0.0001.

In contrast to many other model organisms, *C. elegans* possesses neither DMC1 nor γH2AX, and therefore levels of DSBs were quantified by counting the number of RAD-51 per oocyte precursor cell. Meiotic nuclei move unidirectionally through the *C. elegans* gonad as oocyte precursor cells mature, and so we divided the whole gonad into seven equal-length zones and scored the number of RAD-51 per nucleus in each zone to assess timing and levels of DSBs during meiotic prophase. Based on RAD-51 immunofluorescence, both *spo-11(3A)* and *spo-11(ΔC39)* mutant oocytes showed profoundly reduced DSB levels while the *spo-11(ΔC21)* mutant showed reduced DSB levels to a lesser extent in zone 5 (**Figure 4A, B**).

Previous studies in *C. elegans* have shown that meiotic checkpoints monitor the formation of CO intermediates (39–41), extending the time spent in the leptotene/zygotene transition zone and early pachytene for nuclei lacking CO intermediates. The delayed RAD-51 peak in the *spo-11(ΔC39)* mutant oocytes likely reflects activation of a meiotic checkpoint extending the DSB production window in this mutant. These phenotypes support the idea that the SPO-11 N-terminus and C-terminus are both important to interact with DSB-1/2 at the predicted surfaces.

In order to assess SPO-11 protein levels in these mutants, we used CRISPR to insert a 3xFLAG-tag near the C-terminus of the endogenous *spo-11* gene, and regenerated our *spo-11* mutants in this tagged gene (**Supplemental Figure S4A, S4B**). The *spo-11(wt)::3xFLAG* strain showed a comparable embryonic viability to the N2 wild type, and thus the intragenic FLAG tagging did not interfere with SPO-11 functions (embryonic viability: N2 wild type 99.9% n=3 of P0, *spo-11::3xFLAG* 99.8% n=9 of P0 animals, **Supplemental Figure S4A**). Western blotting using FLAG antibodies showed that *spo-11(3A)::3xFLAG* and *spo-11(ΔC21)::3xFLAG* mutants expressed SPO-11 at the levels comparable to the *spo-11(wt)::3xFLAG* control whereas *spo-11(ΔC39)::3xFLAG* mutants had a 58% reduction in SPO-11 abundance, raising the possibility that the SPO-11 C-terminus contributes to protein stability (**Supplemental Figure S4B**). Although SPO-11 protein levels were reduced in the *spo-11(ΔC39)::3xFLAG* mutants, the reduced DSB production seen in this mutant is likely due to the missing SPO-11 C-terminal region and not to reduced protein amount, since worms heterozygous for a *spo-11* null deletion (*spo-11(ok79)/+* heterozygous) are known to be capable of generating sufficient DSBs and crossovers (42).

Next, to examine the rate of crossover formation in these mutants, we scored the number of DAPI-stained bodies in mature oocytes at diakinesis. In the wild type (2n=12), mature oocytes have 6 DAPI-stained bodies corresponding to 6 chiasma-linked homologous chromosomes. Consistent with reduced DSB levels, univalent chromosomes were detected in both *spo-11(3A)* and *spo-11(ΔC39)* mutants, whereas the *spo-11(ΔC21)* mutant showed normal levels of 6 bivalents (**Figure 4C**). This suggests that although the *spo-11(ΔC21)* mutant had reduced levels of DSBs, it generated sufficient numbers of DSBs to form one crossover per chromosome. During oogenesis in *C. elegans*, it has been shown that a meiotic cell cycle checkpoint induces apoptosis in oocyte precursor cells which fail to generate a crossover on every pair of homologous chromosomes, and the nuclei without crossovers are preferentially culled during meiotic prophase (43). This crossover checkpoint mechanism may remove oocyte precursor cells carrying an insufficient number of crossovers, if there are any, in the *spo-11(ΔC21)* mutant.

Consistent with crossover formation rates, embryonic viability was reduced in *spo-11(3A)* and *spo-11(ΔC39)* mutants whereas the *spo-11(ΔC21)* mutant showed normal levels of embryonic viability, comparable to the wild type (**Figure 4D**). These effects of SPO-11 mutations to the predicted interaction surfaces show functional importance of these regions.

### Mutations in *dsb-1* at the DSB-1-SPO-11 interaction surface lead to reduction in DSB formation *in vivo*

To further examine the SPO-11 and DSB-1 interaction at the YISETPVK motif, we attempted to rescue the *spo-11(3A)* mutation by introducing complementary mutations to the interacting DSB-1 surface. We used Rosetta (44) to redesign DSB-1 and generated the *dsb-1(RD: <u>r</u>e<u>d</u>esigned)* mutant by introducing 10 point mutations (S97K, V99T, R100I, K101L, Y104F, S106T, T108K, V110K, K111L, R116T) around the conserved YISETPVK motif, attempting to restore the DSB-1 binding to SPO-11(3A) protein. However, the *dsb-1(RD)* mutation did not restore *spo-11(3A)* mutant’s embryonic inviability (**Supplemental Figure S5A**). In fact, the *dsb-1(RD)* mutant by itself abrogated DSB formation and crossover formation, and showed very low embryonic viability, indicating that the conserved YISETPVK sequence and its structure is important for DSB formation in the wild type (**Figure 5A**,**B**, **Supplemental Figure S5A**). We also introduced these mutations in *3xFLAG*::*dsb-1* at the endogenous site, and found that DSB-1^RD^ protein levels were reduced to roughly half of that in the *3xFLAG*::*dsb-1(wt)* control (**Supplemental Figure S5B**). Although DSB-1 protein levels were reduced in the *dsb-1(RD)* mutant, the near-complete loss of DSBs in this mutant is likely due to the point mutations at the YISETPVK motif disrupting the interaction with SPO-11, but not due to reduced protein levels. This is because a similar reduction of DSB-1 in worms heterozygous for a *dsb-1* null mutation (*dsb-1(we11)/*+ heterozygous mutant) does not abrogate DSB formation (**Supplemental Figure S5C, S5D**). These data indicate that failure in DSB formation in the *dsb-1(RD)* mutant is most likely due to the point mutations in and around the YISETPVK motif disrupting interaction with SPO-11.

**Figure 5:**
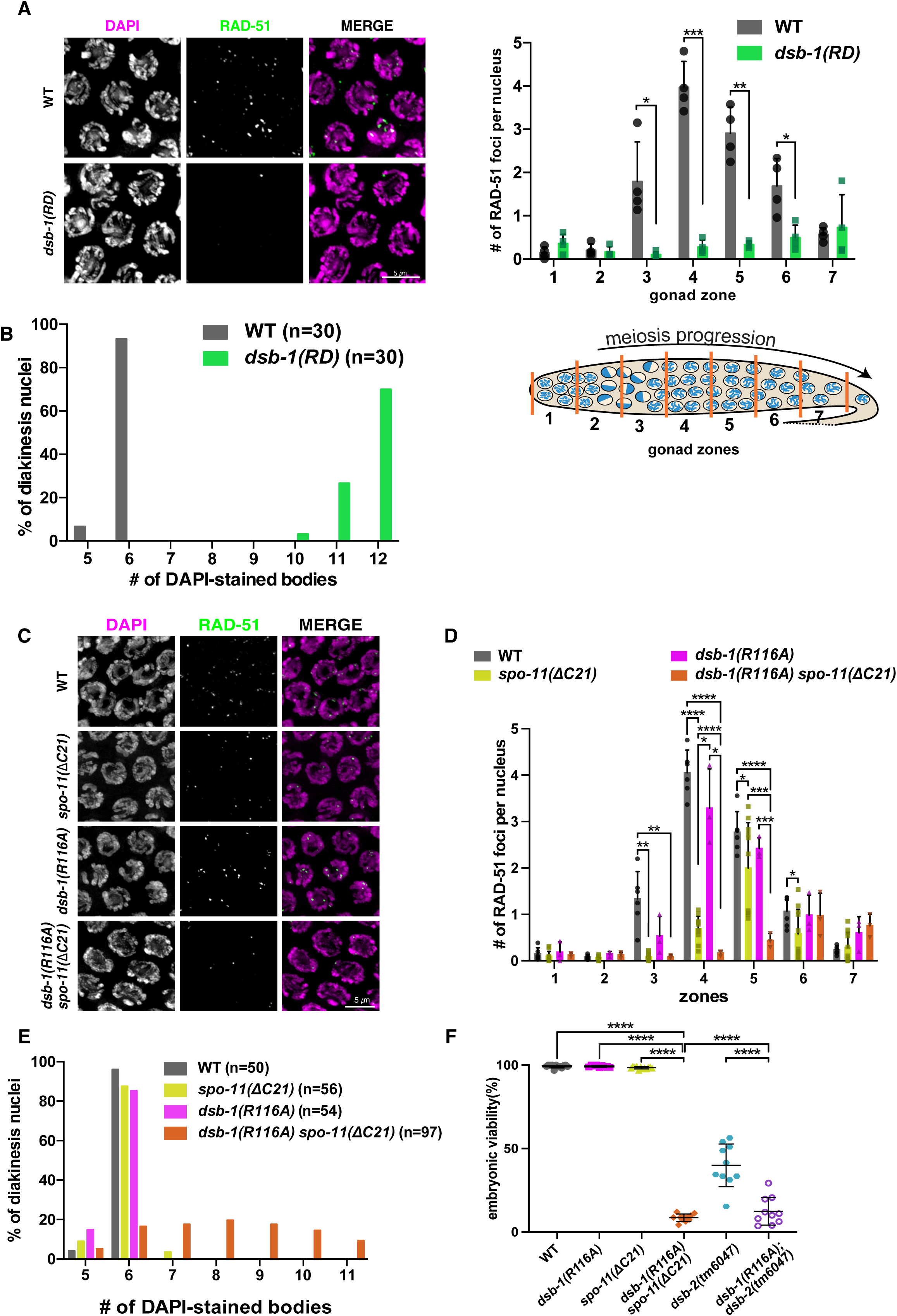
DSB-1 mutants at the SPO-11 interaction surface reduced DSB production in oocytes (A) (Left) Representative immunofluorescence images of RAD-51 foci in mid-pachytene oocytes (zone 4) of the gonads for each genotype indicated. Scale bar, 5 µm. (Right) Quantification of RAD-51 foci per oocyte nucleus in the gonads of indicated genotypes on the left. Data are presented as mean±SD (bar, overall mean; dot, mean of each gonad). The numbers of scored nuclei in zone 1-7 were as follows: for the wild type (4 gonads), 132, 195, 214, 202, 193, 130, 62; for *dsb-1(RD)* (4 gonads), 187, 269, 272, 245, 205, 202, 104. Statistical significance was assessed using a two-tailed t-test with Welch’s correction, **p<0.01, ***p<0.001. **(B)** Distribution of DAPI-stained bodies in diakinesis oocytes, shown as percentages, for the indicated genotypes in **(A)**. **(C)** Representative immunofluorescence images of RAD-51 foci in mid-pachytene oocytes (zone 4) of the gonads for each genotype. Scale bar, 5 µm. **(D)** Quantification of RAD-51 foci in the gonads of indicated genotypes in **(C)**. Data are presented as mean±SD (bar, overall mean; dot, mean of each individual gonad; The numbers of scored nuclei in zone 1-7 were as follows: for the wild type (6 gonads), 205, 285, 291, 366, 321, 241, 105; for *spo-11(ΔC21)* (11 gonads), 646, 850, 783, 891, 732, 521, 140; for *dsb-1(R116A)* (3 gonads), 100, 139, 131, 166, 164, 145, 61; for *dsb-1(R116A) spo-11(ΔC21)* (3 gonads), 97, 140, 154, 190, 157, 108, 28. Statistical significance was assessed using a two-tailed t-test with Welch’s correction, **p<0.01, ***p<0.001, ****p<0.0001. **(E)** Distribution of DAPI-stained bodies in diakinesis oocytes, shown as percentages, for the indicated genotypes in **(C)**. **(F)** Embryonic viability of self-fertilized progeny from hermaphrodites of the indicated genotypes. Data are presented as percentages. Statistical significance was assessed using a two-tailed t-test with Welch’s correction, ****p<0.0001.

As we analyzed the *3xFLAG::dsb-1* strain, we noted that the 3xFLAG-tag moderately reduced DSB production (**Supplemental Figure S6A**). However, 3x*FLAG::dsb-1(wt)* hermaphrodite worms showed normal embryonic viability (99.1%, n=10 P0) at the standard culturing temperature (20 degree) comparable to the wild type (99.3%, n=3 P0), suggesting the effect of FLAG tag on DSB-1 activity is rather minimal (**Supplemental Figure S6B**). For this reason, phenotypic analysis of *dsb-1* mutants in this study was limited to those without the FLAG insertion, while FLAG-tagged strains were used to assess protein levels of various *dsb-1* mutants compared to the control strain by Western blotting.

We next decided to examine the predicted *trans*-SPO-11 binding mediated by DSB-1 R116, and generated a *dsb-1(R116A)* mutant converting Arginine 116 to Alanine to disrupt potential interactions. Since the SPO-11 residue M353 that is most often predicted to contact DSB-1 R116 is predicted to be very close to the catalytic tyrosine (Y119), we did not attempt to mutate the SPO-11 side of this predicted interaction. The *dsb-1 (R116A)* mutation by itself only slightly reduced DSB formation (**Figure 5C, 5D**) and showed embryonic viability comparable to the wild type (99.8% embryonic viability in N2 v.s. 99.2% in *dsb-1 (R116A)*, **Figure 5F**). Since the predicted interaction between DSB-1 R116 and SPO-11 represents only one of two possible *trans-*interactions, the other being through the C-terminal tail of SPO-11, we next decided to combine the R116A mutation with the *spo-11(ΔC21)* deletion, which by itself resulted in normal embryonic viability. The double *dsb-1(R116A) spo-11(ΔC21)* mutants showed a striking failure of DSB formation, crossover formation (**Figure 5C, D, E**) with reduced embryonic viability (8.6%) compared to normal viabilities seen in the *spo-11(ΔC21)* single (98.6%) or *dsb-1 (R116A)* (99.2%) mutant (**Figure 5F**). This synergistic effect suggests that the R116A mutation does mildly compromise DSB-1 activity, possibly by weakening the DSB-1/*trans-*SPO-11 interaction, but the defect is compensated for by the SPO-11 C-terminal extension.

In addition, we combined the R116A mutation with a *dsb-2* null mutation, *dsb-2(tm6047)*. In contrast to DSB-1 being absolutely essential for SPO-11 activity, DSB-2 is not strictly required for gamete production especially in young hermaphrodites. Null mutants of *dsb-2* show 40.2% embryonic viability in self-fertilized progeny of hermaphrodites, and show reduced stability of DSB-1 protein as well as reduced DSB levels in oocytes (**Figure 5F, 6A**) (35–37). The *dsb-1(R116A); dsb-2(tm6047)* double mutants showed a further decrease to 12% embryonic viability, compared to 99.2% for *dsb-1(R116A)* single mutants, showing the *dsb-1(R116A)* mutation also exacerbates the reduction in DSB formation caused by loss of DSB-2. In order to verify that the R116A mutation did not alter DSB-1 protein stability, we introduced this mutation in 3xFLAG-tagged *dsb-1* at the endogenous site and found that DSB-1^R116A^ protein was expressed at similar levels compared to the control (**Supplemental Figure S6C**). Taken together, these observations support the idea that DSB-1/SPO-11 interaction is stabilized by the predicted interaction at the R116 residue of DSB-1, and suggest that at least one *trans*-SPO-11 interaction with DSB-1 or DSB-2 is required for DSB formation.

### The evolutionarily coemerged *dsb-2* and *spo-11* C-terminus ensure robust DSB formation in oogenesis

Our phylogenetic tree-based MSA analysis showed coemergence of DSB-2 and the SPO-11 C-terminus extension during *Caenorhabditis* speciation, hinting at a possible interaction between them. By AF modeling, SPO-11 C-terminal fragment in isolation was predicted to bind to the isolated PH domains of both DSB-1 or DSB-2 (**Figure 2E**) and our pull-down assay showed the SPO-11 C-terminus can bind to DSB-1 or DSB-2’s PH domain, but these showed no binding preference for DSB-2 over DSB-1. On the other hand, AF modeling with the complement ([DSB-1-2-3]×2, SPO-11×2, DNA) consistently predicted structures with the SPO-11 C-terminus bound to DSB-2 rather than DSB-1 (**Figure 2D**). We therefore decided to assess phenotypic similarity between our *spo-11(ΔC39)* mutants and *dsb-2* mutants. In contrast to *spo-11* or *dsb-1* null mutations which completely abrogate DSBs, *dsb-2* hermaphrodite mutants are competent for low levels of DSB formation (45). We found that sperm viability is mostly normal in *dsb-2* males (98.3% for wild type sperm viability v.s. 91.4% in *dsb-2(me96)* when crossed to control *fog-2* females (obligate females produced by the *fog-2* mutation)) whereas embryonic viability of the self-fertilized progeny of *dsb-2* hermaphrodites producing both sperm and oocytes is 45.4%, suggesting that the net oocyte viability is around 49.7%. We found that *spo-11(ΔC39)* males also produced viable sperm comparable to wild type males (99.3% in wild type sperm v.s. 97.4% in *spo-11(ΔC39)* sperm), whereas viability of the self progeny of *spo-11(ΔC39)* hermaphrodites was 26.8%, suggesting that oocyte viability is 27.5% (**Figure 6A**). These data raised the possibility that DSB-2 and SPO-11 C terminal extension co-evolved to function mainly in oogenesis but not in spermatogenesis. To understand the sexually dimorphic effect of these mutants, we scored the number of RAD-51 foci in spermatocytes in males and compared this to the number in oocytes. We found reduction in DSB levels in both oocytes and male spermatocytes in both *spo-11(ΔC39)* or *dsb-2* null mutants compared to wild type males or oocytes. However, the loss of DSBs was much more pronounced in oocytes (6.6% (*spo-11(ΔC39)*) or 5.8% (*dsb-2*) of WT levels) compared to spermatocytes (50% (*spo-11(ΔC39)*) or 29% (*dsb-2*) of WT levels) at the peak zone (mean number of RAD-51 foci per nucleus at peak zone 4: wt oocytes = 3.8 (n=8) v.s. wt spermatocytes =6.9 (n=3), *ΔC39* oocytes =0.25 (n=8), *ΔC39* spermatocytes = 3.5 (n=3), *dsb-2* null oocytes =0.22 (n=2), *dsb-2* spermatocytes = 2.0 (n=3)). Additionally, oocytes showed an overall lower number of DSBs compared to spermatocytes in all genotypes (**Figure 6B**: RAD-51 number in *ΔC39* oocytes is the same data presented in **Figure 4B**, shown here for visual comparison to spermatocytes). Recently, a similar result was reported for *dsb-2* mutant spermatocytes compared to wild type spermatocytes (45). Previous studies have shown that the kinetics of DSB formation are different between spermatogenesis and oogenesis: DSBs are generated earlier in meiotic prophase and they reach higher numbers in spermatogenesis compared to oogenesis in *C. elegans.* (45). Although the *dsb-2* null and *spo-11(ΔC39)* mutations reduced DSB levels in both spermatocytes and oocytes, elevated activity in DSB formation during spermatogenesis likely secured a sufficient number of DSBs for viable sperm production in these mutants. Male meiosis in *C. elegans* has also been reported to distributively segregate achiasmate chromosomes to opposite spindle poles in roughly equal mass at meiosis I more efficiently than oocytes presumably due to differential organization of meiotic spindles (46). Indeed, *spo-11* or *dsb-1* null mutant sperm have 15-18% of viability when fertilizing *fog-2* control oocytes (99.3% for wt sperm: n= 16, 18.2% for *spo-11(me44)* sperm: n=12, 15.2% for *dsb-1(we11)* sperm: n=10) whereas the self progeny of *spo-11* or *dsb-1* null hermaphrodites are known to have close to zero viability, suggesting that their oocytes have nearly zero viability (**Figure 6A**)(42). The combined evidence suggests that elevated intrinsic DSB production activity, less reliance on the SPO-11 C-terminus and on DSB-2, and more efficient achiasmate segregation mechanisms during spermatogenesis could together explain the nearly intact sperm viability in *dsb-2* null and *spo-11(ΔC39)* mutants.

**Figure 6:**
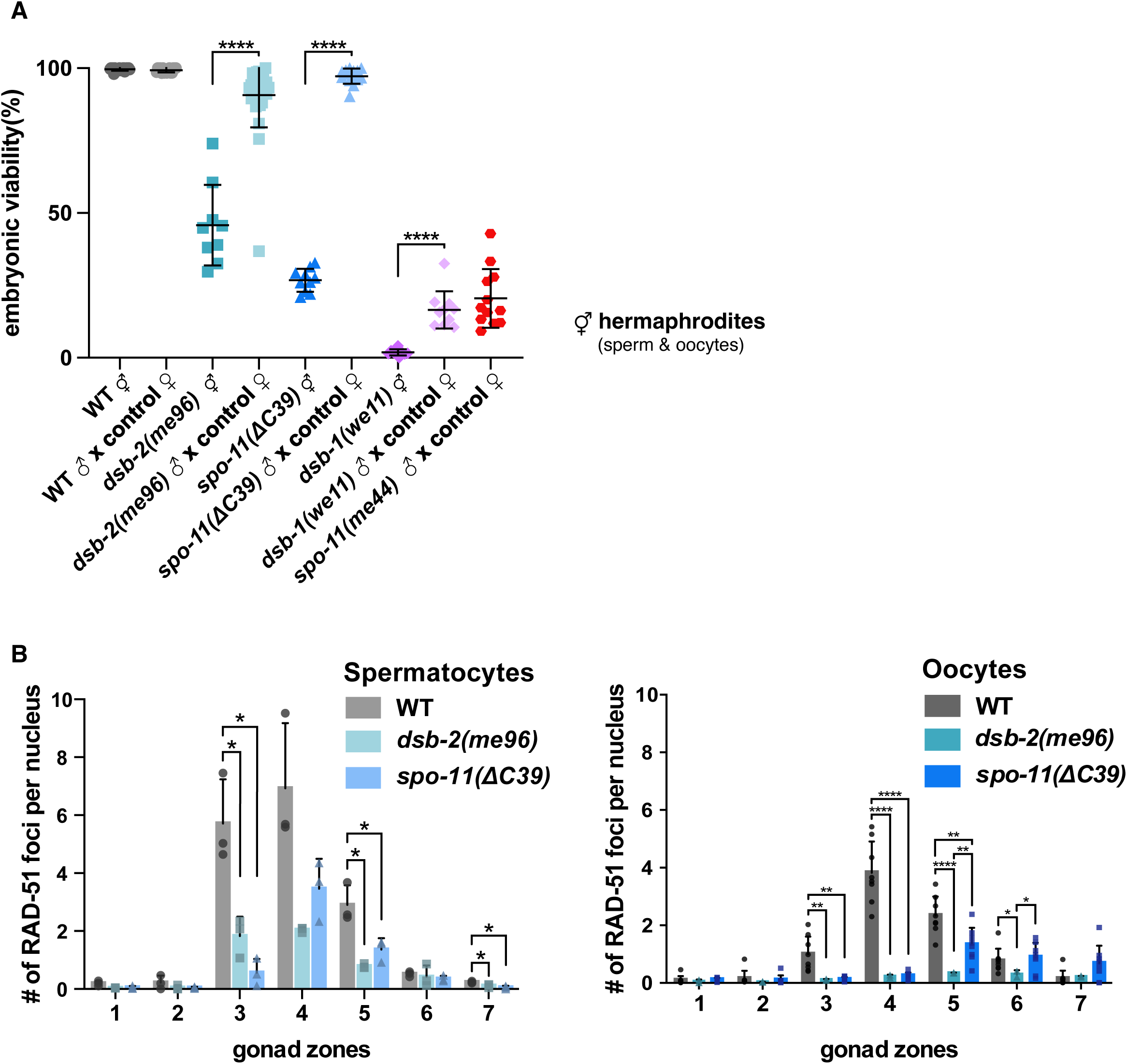
Reduced DSB formation preferentially impairs oogenesis in *dsb-2(me96)* and *spo-11(ΔC39)* mutants. **(A)** Embryonic viability of self-progeny from hermaphrodites of the indicated genotypes (wild type, *dsb-2(me96)*, *spo-11(ΔC39)*, and *dsb-1(we11)*), reflecting the combined contributions of oocytes and sperm. Embryonic viability of cross-progeny from males of the indicated genotypes crossed with *fog-2(q71)* females, reflecting sperm viability in the wild type, *dsb-2(me96)*, *spo-11(ΔC39)*, *spo-11(me44)*, and *dsb-1(we11)* mutants. Data are presented as percentages. Statistical significance was assessed using a two-tailed t-test with Welch’s correction, ****p<0.0001. The embryonic viability of *ΔC39* self progeny is the identical data presented in Figure 4D and is shown here for a comparison to the male viability **(B)** Quantification of RAD-51 foci in the male gonads (left) or hermaphrodite gonads going through oogenesis (right) in the wild type, *dsb-2(me96)* and *spo-11(ΔC39)* mutants. Data are presented as mean±SD (bar, overall mean; dot, mean of each individual gonad; The numbers of scored nuclei in zone 1-7 were as follows: for wild type male (3 gonads), 107, 102, 100, 79, 64, 81, 77; for *dsb-2(me96)* male (3 gonads), 104, 100, 115, 100, 68, 75, 70; for *spo-11(ΔC39)* male (3 gonads), 150, 142, 132, 117, 85, 101, 92; for wild type hermaphrodite going through oogenesis (8 gonads), 549, 654, 663, 730, 571, 471, 233; for *dsb-2(me96)* hermaphrodite (2 gonads), 57, 91, 90, 102, 73, 71, 43; for *spo-11(ΔC39)* hermaphrodite (8 gonads), 397, 586, 589, 610, 553, 387, 120. Statistical significance was assessed using a two-tailed t-test with Welch’s correction, **p<0.01, ****p<0.0001.

Previous studies have shown that the quality of oocytes worsened with age in *dsb-2* mutant hermaphrodites (35, 37). Embryos produced by self-fertilization of young (1 day post L4 stage) *dsb-2* hermaphrodites are 69.3% viable (n=9) whereas embryos derived from older *dsb-2* hermaphrodites (2 days post L4 stage) are 46.8% viable (n=9) and 25.8% viable (n=9) for further older *dsb-2* hermaphrodites (3 days post L4 stage) due to decreasing efficiency of crossover formation with age. Since hermaphrodites first produce sperm during the last larval stage, and switch to oogenesis at the adult stage, this age effect of hermaphrodite adults is attributed to the worsening quality of oocytes. Although hermaphrodites produce sperm only at the last larval stage before adulthood, male worms can produce sperm throughout their adult life. We therefore wondered if there is any age effect on the quality of sperm in *dsb-2* males, and compared viability of sperm produced by males at different ages by crossing them to *fog-2* obligate females. We did not find any age effect on the quality of sperm from *dsb-2* males (sperm viability of day 1 *dsb-2* adult males 93.7% (n=10), day 2 adult males 87.0%(n=8), day 3 adult males 93.6%(n=10), day 4 adult males 91.0% (n=4)). This suggests that an unknown factor changes with age specifically in oocytes leading to gradual DSB reduction during oogenesis but not in spermatogenesis in the absence of DSB-2.

Although *dsb-2* null and *spo-11(ΔC39)* mutants reduced DSB production to similar levels, we noted phenotypic differences in these mutants. Previous studies have shown that DSB-1 protein levels are reduced in *dsb-2* null mutants, indicating that DSB-2 stabilizes DSB-1 (35, 36). In contrast, we found that DSB-1 protein levels are not reduced in *spo-11(ΔC39)* mutants, indicating that DSB-2’s role in DSB-1 stability is independent from its possible interaction with the SPO-11 C-terminus (**Supplemental Figure S6D**). This suggests the possibility that DSB-2 may stabilize DSB-1 in the DSB-1-2-3 trimer complex independently of SPO-11. In addition, as mentioned above, the embryonic viability of self-fertilized progeny in *dsb-2* null mutants declined with age (day 1, 69.3%; day 3, 25.8%)(35), whereas embryonic viability in *spo-11(ΔC39)* mutants did not decrease with age, but rather showed a modest increase (**Supplemental Figure S6E**). These differences suggest that the extended SPO-11 C-terminus and/or DSB-2 do not strictly work together but have at least some roles that are independent from each other. Taken together, our data suggest that DSB-2 and the SPO-11 C-terminal extension region contribute to robust DSB production both in spermatogenesis and oogenesis, but their roles are more critical for oogenesis.

## Discussion

In this study, by taking advantage of comparative genomics in *C. elegans* and related species, we surveyed evolutionary correlation and conservation between functionally related proteins required for meiotic DSB initiation. We identified a key protein domain in DSB-1 that, by binding to SPO-11, likely permitted the loss of TOPOVIBL during *Caenorhabditis* speciation. Our biochemical and *in vivo* mutagenesis data suggest that, in *C. elegans*, DSB-1-2-3 proteins can directly bind to SPO-11 at several surfaces, and that one DSB-1 can bind to two SPO-11 molecules simultaneously, thereby promoting SPO-11 dimerization.

AlphaFold3 prediction and mutant phenotypes identified three putative interactions between DSB-1 and SPO-11: (1) the YISETPVK motif of DSB-1 with the RKIEFALAD motif of *cis-*SPO-11; (2) R116 of DSB-1 with *trans-*SPO-11 residues, especially M353; and (3) the acidic C-terminus of SPO-11 (*cis-*and *trans-*not distinguished) binding to basic residues in the PH domain of DSB-1 and DSB-2. Our genetic analysis of the mutants *spo-11(3A)*, *dsb-1(RD)*, and combinations of *dsb-1(R116A)* and *spo-11(ΔC21)* shows that DSB formation absolutely requires the first of these interactions, and at least one of the latter two. From this evidence, we conclude that each DSB-1 molecule likely engages in one *cis-*and one *trans-*interaction to promote SPO-11 dimer formation and thus DNA breaks.

Although a previous study surveying 11 *Caenorhabditis* species had pointed out that acquisition of the SPO-11 C-terminal extension is found in *Caenorhabditis* species without TOPIVBL, the further analysis we performed here shows that many species outside of the Elegans and Japonica groups diverged after the loss of TOPOVIBL, but before emergence of the C-terminal extension. Our analysis of 54 *Caenorhabditis* species shows instead that this extended C-terminus of SPO-11 is found specifically in the Elegans and Japonica group species that diverged after an ancient duplication of *dsb-1* (emergence of *dsb-2*).

This correlation, as well as the higher tendency to see DSB-2 association with SPO-11 C-termini in AF models, raised the possibility that the SPO-11 C-terminus extension region interacts with DSB-2 more stably than with DSB-1 *in vivo*. However, we currently have no evidence for this, as our pulldown assay shows that the DSB-1 and DSB-2 PH domains both bind to the SPO-11 C-terminus extension. Further, *dsb-2* has been lost twice in lineages that once possessed it: once in the ancestor of the Elegans group sub-clade containing *C. brenneri, C. sp44, C. sp48,* and *C. sp51*, and once in *C. panamensis*, a Japonica group species, and these species have all maintained their SPO-11 C-terminus extension (**Supplemental Figure S2**). Further analysis will be needed to understand the evolutionary process of acquisition of DSB-2 and SPO-11 C-terminus extension in the nematode lineage. There are currently ∼30 diverse *Caenorhabditis* species with unsequenced or incomplete genomes, and these may shed further light on these relationships.

Instead of the SPO-11 C-terminal extension, what does correlate with loss of TOPOVIBL is the SPO-11-interacting [YF]ISE[ST]PXK motif of DSB-1. Although *dsb-1* and *dsb-2* are paralogs, the [YF]ISE[ST]PXK motif has been retained only in one paralog, represented by *C. elegans dsb-1*. The other paralog has lost the motif and became the markedly diverged *dsb-2* in the Elegans group, while in the Japonica group it has persisted as a less diverged paralog (**Supplemental Figure S2**). Species outside of the Elegans and Japonica groups lack these ancient paralogous pairs, but do present 6 cases of more recent (i.e., terminal or near-terminal) duplication of *dsb-1*, which may reflect ongoing diversification.

Examination of the phylogenetic tree (**Figure 1**) presents one apparent species exception, *C. plicata,* to the correlation between loss of TOPOVIBL and gain of the [YF]ISE[ST]PXK motif, since neither the motif nor TOPOVIBL can be detected. However, since DSB-1 in its sister species *C. bovis* does have the [YF]ISE[ST]PXK motif, the common ancestor of *C. bovis* and *C. plicata* likely possessed the motif, which subsequently was lost in *C. plicata*. One possibility is that the [YF]ISE[ST]PXK-containing *dsb-1* emerged as a duplication of the ancestral Rec114, and this did not persist in *C. plicata*.

The significance of the simultaneous duplication of *dsb-1* (emergence of *dsb-2*) and emergence of the SPO-11 C-terminal extension is not clear. DSB-2 is known to stabilize DSB-1 and ensures DSB production in oocytes in older hermaphrodites. We show here that both of these innovations are more important for DSB production in oogenesis than in spermatogenesis. Our previous work also suggested that the activity of DSB-1, but not DSB-2, is regulated by phosphorylation by ATR kinase, presumably to prevent excessive DSB production (35). In addition, both DSB-1 and DSB-2 are actively degraded by an unknown mechanism upon satisfaction of a crossover-ensuring cell cycle checkpoint. (36, 37). Since having too many DSBs leads to genomic instability, while too few DSBs lead to lack of crossovers and chromosome missegregation during meiosis, adjusting the activity and protein levels of DSB-1 is likely very important for gametogenesis. *Caenorhabditis* species include both gonochoristic (male/female) and androdioecious (male/hermaphrodite) worms and have diverse lifestyles, durations of reproduction and rates of oocytes/sperm production. Innovations such as DSB-2 and the SPO-11 C-terminal extension may have been selected to fine-tune or extend DSB-1 lifetime specifically to ensure the quality of oocytes as worms alter the mode of reproduction during the course of evolution.

The breadth and availability of nematode genome and proteome sequence data let us identify co-evolution of functionally related protein domains in this study. Increasing efforts in whole-genome sequencing of more species may empower comparative genome studies to identify functional relationships between relevant proteins or protein domains similar to this study. Our results have shown that multifaceted, direct interactions between DSB-1/2 and SPO-11 replaced TOPOVIBL during speciation, and suggest that one DSB-1 binds to two SPO-11 molecules simultaneously to promote their dimerization, enabling programmed DSBs in meiotic prophase.

## Materials and methods

### *C. elegans* strains and antibodies

All *C. elegans* strains in this study were maintained on nematode growth media (NGM) plates following standard conditions at 15∼25°C (47). The N2 Bristol strain was used as the wild-type. Adult homozygous mutants derived from balanced, heterozygous mutants were used for cytological and genetic experiments. A list of all strains and antibodies used is provided in the **Supplemental Table S1**.

### Generation of bacterial constructs for pull-down assay

#### MBP fused protein

The pGEX-6p-1-MBP construct was derived from the pGEX-6p-1 vector. The pGEX-6p-1 vector was linearized by PCR amplification and MBP cDNA sequence was amplified by PCR. The two PCR products were ligated using SLiCE assembly (48) to create a pGEX-6p-1-MBP empty vector. This empty vector was linearized by PCR amplification, and DNA regions coding for SPO-11 (31-150aa) and SPO-11 (405-425aa) were amplified by PCR using a *C. elegans* cDNA library. The amplified backbone vector was combined with each amplified cDNA fragments using SLiCE assembly to create pGEX-6p-1-MBP-SPO-11 (31-150aa) and pGEX-6p-1-MBP-SPO-11 (405-425aa) constructs.

### 10xHis tag fused protein

The pET-16b vector was linearized by PCR amplification, and DSB-1 (1-112aa) and DSB-2 (1-108aa) were amplified by PCR using *C. elegans* cDNA library. The linearized pET-16b vector was combined with the amplified cDNA fragments respectively to create pET-16b-DSB-1 (1-112aa) and pET-16b-DSB-2 (1-108aa) constructs using Gibson assembly (New England Biolabs, E2611L). The PCR primers used for cloning are shown in **Supplemental Table S**2.

### Immunofluorescence and imaging

Immunostaining was performed as described in Rillo-Bohn et al. (49) modified thus: homemade mounting medium (250 mM Tris, 1.8% w/v *n*-propyl gallate in glycerol) and Matsunami No. 1S (Matsunami, Osaka, Japan; nominal width 0.17mm) coverslips were used. Young adult hermophrodites producing oocytes (1 day post-L4 larval stage) or young adult males producing sperm were used for immunostaining. A list of antibodies used is provided in **Supplemental Table S3**. Images of the gonad regions at the coverslip-proximal side were captured by a Deltavision personalDV microscope (Applied Precision) equipped with a CoolSNAP ES2 camera (Photometrics) at a room temperature of 20–22°C, using a 100× UPlanSApo 1.4NA oil immersion objective (Olympus, Tokyo, Japan) and immersion oil (LaserLiquid; Cargille, Cedar Grove, NJ) at a refractive index of 1.513. The Z spacing was 0.2 µm and raw images were subjected to constrained iterative deconvolution followed by sub-pixel Z shifts to correct chromatic aberration. Image acquisition and deconvolution was performed with the softWoRx suite (Applied Precision/GE Healthcare, Chicago, IL). Image postprocessing for publication was limited to linear intensity scaling and maximum-intensity projection using OMERO (Burel et al., 2015). For RAD-51 quantification, we used *ndevio* (https://github.com/ndev-kit/ndevio), *napari* (https://napari.org) (50), and *multiview-stitcher* (51) to read, stitch and display the gonads before manually scoring the foci of genotype and meiotic-stage nuclei using a napari plugin we developed (https://github.com/carltonlab/carltonlab-napari-tools). Using this plugin, focus scoring is performed on isolated, genotype-blinded nuclei. Statistical comparisons of the mean number of RAD-51 per nucleus between equal zones were performed via two-tailed t test with Welch’s correction.

For bivalent/univalent chromosome counting, completely resolvable contiguous DAPI-positive bodies were counted in 3D stacks as described previously (52). With this criterion, chromosomes that happen to be touching can occasionally be counted as a single DAPI body, resulting in a small number of counts less than 6.

### Embryonic viability quantification

To score embryonic viability and male progeny derived from self-fertilization of hermaphrodites, L4 larval stage worms (P0s) from each genotype were picked individually onto plates and transferred to fresh plates every 24 hr for 4–5 days to score the whole brood. Unhatched eggs remaining on the plates 48 hr after being laid were counted as dead eggs. Viable F1 progeny and males were scored 72 hr after P0s were transferred to new plates. Embryonic viability for each day was calculated separately from the daily brood counts. Day 1 and day 3 embryonic viability within each genotype were compared using a one-tailed *t*-test with Welch’s correction.

To score sperm viability, one L4 stage *fog-2(q71)* female was placed on a single plate with 4 mixed stage adult males for 40 to 48 hours and allowed to mate and lay eggs, after which all the P0 parents were removed. 24 hours after removal of parents, unhatched eggs were scored as dead eggs; and 48 hours after removal of parents, hatched progeny were scored as viable.

To compare sperm viability derived from *dsb-2(me96)* males at different ages, first *dsb-2(me96)* mutants including both hermaphrodites and males were synchronized at the L1 larval stage by bleaching and starvation, and released on NGM plates with OP50 bacteria as a food source. Males were picked for mating with *fog-2(q71)* obligate females either after 56h post L1 release (as day 1 adult males), 80h post L1 (day 2 adult males), 104h post L1 (day 3 adult males) or 128h post L1 (day 4 adult males), and embryonic viability of their cross progeny laid by each *fog-2(q71)* female was scored as in the above assay of sperm viability counting. Mating plates yielding no progeny were excluded due to either failure of mating or shortage of sperm in males. Until being picked for mating, *dsb-2* males were kept on a plate with *dsb-2* hermaphrodites, to prevent sperm hoarding until further mating with *fog-2* females. Since no age effect was found in sperm viability derived from *dsb-2(me96)* males, all embryonic viability data from males from all ages were pooled to generate the data points in **Figure 6A**.

### AlphaFold structure prediction

Predictions were generated using the AlphaFold3 server (alphafoldserver.com) with new random seed assignments for each repeated prediction. Raw .cif files from AlphaFold3 were first regularized using Phenix (53) geometry minimization (version 1.21.2-5419) to eliminate clashes. Subsequently, structures were visualized and superimposed with ChimeraX (54), and contacts visualized using the "circoscontacts" plugin (https://github.com/carltonlab/circoscontacts). DNA in all models was composed of one 60-base ssDNA chain derived from *C. elegans* chromosome III with genome-representative GC content, and two 30-base chains, each complementary to a contiguous half of the 60-base chain, an arrangement that AlphaFold3 consistently folds into a nicked dsDNA structure.

### Multiple sequence alignment, orthology clustering and gene tree inference

We obtained protein, genome, and annotation files for 54 *Caenorhabditis* species from WormBase ParaSite (Howe et al., 2017) and the Caenorhabditis Genomes Project website (formerly caenorhabditis.org), presently archived at https://zenodo.org/records/12633738 (55). Out of these 54 species, 42 possessed annotated, unambiguous DSB-1 proteins and 50 possessed annotated SPO-11 proteins; unannotated sequences were recovered manually from the genomes using query sequences from neighboring species. For proteomes lacking annotated genes, miniprot 0.18 in protein2genome mode was used to generate plausible gene models using phylogenetically close protein sequences as queries. Genomic similarity searches were performed with TBLASTN (BLAST+ 2.12.0+), using both standard settings and more sensitive searches with BLOSUM45 and reduced word size to detect weak homologous segments independent of gene prediction. Then, all manually-recovered and previously-annotated proteins were reconfirmed with miniprot. In two cases, *C. tropicalis* and *C. bovis*, the C-terminal region of DSB-1 could not be unambiguously determined from the current genomes; however, the N-terminal region containing the [FY]ISE[ST]PXK motif and extending into the IDR was present in both. Clustering and gene tree building was performed based on (35): we aligned sequences of the SPO-11 and DSB-1/2 orthogroup sequences using mafft 7.490 (56) and built gene trees using IQ-TREE v3.1.2 (57) under the Q.INSECT+I+R5 substitution model (for SPO-11) or JTT+F+R5 model (for DSB-1/2). A rooted species tree for the 54 *Caenorhabditis* species analyzed was generated with Orthofinder 3.1.1 (58, 59).

We visualized gene trees using iTOL (60) and compared trees side-by-side with MSAs and motifs using msafara (https://github.com/carltonlab/msafara).

### MBP fused Protein purification

All three bacterial constructs expressing MBP or MBP fused protein were transformed into *Rosetta(DE3) pLysS E. coli*. Cells transformed with MBP-SPO-11(31-150aa) were grown in LB medium containing the appropriate antibiotics at 37°C until they reached an OD_600_ of 0.6, followed by induction with 0.5 mM IPTG at 23°C for 4 h. Bacterial pellets were resuspended in column buffer (20 mM Tris-HCl, 200 mM NaCl, 1 mM EDTA, 1 mM DTT, protease inhibitor cocktail), snap-frozen in liquid nitrogen, and sonicated. After centrifugation at 15000 g for 30 min, amylose beads (NEB Inc., E8021S, Ipswich, MA), 20 mM DNaseI and 10 mM MgCl_2_ were added to supernatants and incubated at 4°C overnight with rotation. The mixture was loaded into a column (BIO-RAD Inc #732101, Hercules, CA), washed 15 times with 10 mL wash buffer (20 mM Tris-HCl, 200 mM NaCl, 1 mM EDTA, 1 mM DTT), and eluted 8 times with 1 mL elution buffer (20 mM Tris-HCl, 200 mM NaCl, 1 mM EDTA, 1 mM DTT, 10 mM maltose). The transformed cells of MBP and MBP-SPO-11(405-425aa) were grown in LB medium containing the appropriate antibiotics at 37°C until they reached an OD_600_ of 0.6, followed by induction with 1 mM IPTG at 37°C for 4 -4.5 h. Bacterial pellets were resuspended in column buffer (20 mM Tris-HCl, 200 mM NaCl, 1 mM EDTA, protease inhibitor cocktail), snap-frozen in liquid nitrogen, and sonicated. After centrifugation at 10000 g for 25 min, amylose beads were added to supernatants and incubated at 4°C overnight with rotation. The supernatant was loaded into columns and washed 15 times with 10 mL wash buffer (20 mM Tris-HCl, 200 mM NaCl, 1 mM EDTA) and eluted 8 times with 2 mL elution buffer (20 mM Tris-HCl, 200 mM NaCl, 1 mM EDTA, 10 mM maltose). Protein expression was confirmed by running a 10-20% SDS-PAGE gel (Fujifilm Wako Inc 198-15041, Osaka, Japan), followed by staining with Coomassie Brilliant Blue (CBB) staining kit (Nacalai Inc 30035-14, Kyoto, Japan). The purified proteins were dialyzed with 1x PBS at 4°C for 48 h. Protein concentrations were determined using the Bradford Protein Assay Kit (Thermo Scientific 23200, Waltham, MA).

### 10xHis tag fused protein purification

pET-16b-DSB-1(1-112aa) and pET-16b-DSB-2(1-108aa) were transformed into *Rosetta(DE3) pLysS E. coli*. The transformed cells were grown in LB medium containing the appropriate antibiotics at 37°C until they reached an OD_600_ of 0.6, followed by induction with 0.35 or 0.5 mM IPTG at 18°C overnight or 23°C for 4 h. Bacterial pellets were snap-frozen with liquid nitrogen and resuspended in lysis buffer (50 mM Na_2_PO_4_, 300 mM NaCl, 10 mM imidazole, 0.25% Tween 20, 0.1 mM EGTA, 2 mM β-mercaptoethanol, pH = 8.0) and sonication. After centrifugation at 10000 g for 25 min, the Ni-NTA agarose gel (QIAGEN Inc 30210, Venlo, Netherland), 20 mM DNaseI and 1 mM MgCl_2_ were added into the supernatants and incubated at 4°C overnight with rotation. The mixture was loaded into the column (BIO-RAD #732101, Hercules, CA) and washed 15 times with 10 mL wash buffer (50 mM NaH_2_PO_4_, 300 mM NaCl, 20 mM imidazole, 2 mM β-mercaptoethanol, pH = 8.0) and eluted 10 times with elution buffer (50 mM NaHPO4, 300 mM NaCl, 250 mM imidazole, 2 mM β-mercaptoethanol, pH = 8.0). Protein expression was confirmed by running a 10-20% SDS-PAGE gel (Fujifilm Wako 198-15041, Osaka, Japan), followed by staining with Coomassie Brilliant Blue (CBB) staining kit (Nacalai Inc 30035-14, Kyoto, Japan). The purified proteins were dialyzed with 1x PBS at 4°C for 48 h. Protein concentrations were determined using the Bradford Protein Assay Kit (Thermo Scientific 23200, Waltham, MA).

### Pull-down assay

100 μg of bait protein (MBP fused protein) was incubated with 1:10 molecular ratio of prey protein (10xHis tag fused protein), amylose beads (NEB Inc E8021S, Ipswich, MA) and 1 mM β-mercaptoethanol at 4°C overnight with rotation. The beads were washed 15 times with 1 mL wash buffer (20 mM Tris-HCl, 200 mM NaCl, 1 mM EDTA, 1 mM β-mercaptoethanol) and eluted 5 times with 50 μL elution buffer (20 mM Tris-HCl, 200 mM NaCl, 1 mM EDTA, 10 mM maltose). Protein interaction was confirmed by running a 15-20% SDS-PAGE gel (Fujifilm Wako 198-15301, Osaka, Japan), followed by staining with Coomassie Brilliant Blue (CBB) staining kit (Nacalai Inc 30035-14, Osaka, Japan) and Silver staining kit (Fujifilm Wako 291-50301, Osaka, Japan). Western blot was also performed to confirm the ideal size bands indicating 10xHis tag fusion protein. The 3rd elution was loaded into a 15-20% SDS-PAGE gel (Fujifilm Wako 30035-14, Osaka, Japan), and proteins were transferred to a PVDF membrane (Millipore IPVH00010, Burlington, MA) at 15 V for 40 min. The membrane was blocked with 1% BSA/TBST buffer at room temperature for 1h and probed with primary antibody solution containing 1% BSA at 4°C overnight, followed by washed three times for 10 min, probed with secondary antibody solution containing 1% BSA and washed three times for 10 min. Chemi-Lumi one super (Nacalai Inc 02230-30, Kyoto, Japan) was used to visualize protein bands using an ImageQuant LAS4000 (Cytiva, Marlborough, MA).

### Yeast two-hybrid assay

The pGBT8 vector encoding GAL4 DNA binding domain and the pGAD GH vector encoding GAL4 activation domain were linearized by PCR amplification, and *C. elegans* protein fragments were amplified by PCR using *C. elegans* cDNA library. The linearized vectors and the amplified cDNA fragments were assembled using SLiCE assembly (48). The constructs were transformed into *Saccharomyces cerevisiae* strain HF7c. The transformed yeast cells were grown in the synthetic medium lacking leucine and tryptophan at 30°C until they reached OD_600_ of 0.2. Cells were then serially diluted (10-fold to 10^7^-fold) and spotted on the synthetic medium plates lacking leucine and tryptophan (-L-W), or leucine, tryptophan and histidine (-L-W-H), or leucine, tryptophan and histidine with 2mM 3-amino-1,2,4-trizole (-L-W-H + 2mM 3-AT). Plates were incubated at 30°C for 2 days, and images were acquired using ImageQuant LAS4000 (Cytiva Marlborough, MA).

### CRISPR/Cas9 genome editing

CRISPR/Cas9 using *dpy-10* as a co-CRISPR marker (61, 62) was used to generate *spo-11* and *dsb-1* mutants in this study. For *spo-11::3xFLAG*, *spo-11(3A)*, *spo-11(ΔC21)*, *spo-11(ΔC39)* and *spo-11(3A)::3xFLAG* strain generation, genome editing was done following a previous study ((63): briefly, a 20 µL mixture containing 1.5 µM Cas9 protein, 4.5 µM trans-activating CRISPR RNA (tracrRNA), 4.75 µM CRISPR RNAs (crRNA) targeting *spo-11* (4.4 µM) and *dpy-10* (0.35 µM), 500 ng melted double-stranded DNAs generated from PCR as a repair template and 0.25 µM repair template of *dpy-10* was prepared and injected into gonads of 24 hours post-L4 stage N2 hermaphrodites. For *spo-11(ΔC21)::3xFLAG*, *spo-11(ΔC39)::3xFLAG* strain generation, genome editing was done as follows: 5 µL mixture containing 8.75 µM Cas9 protein, 8.75 µM tracrRNA, 8.75 µM crRNA targeting *dpy-10* (0.67 µM) and *spo-11* or *dsb-1* (8.08 µM), melted double-stranded DNA generated by PCR as a repair template and 0.4 µM repair template of *dpy-10* was injected into gonads of 24 hours post-L4 stage *spo-11(ΔC21)* and *spo-11::3xFLAG* mutants. The concentration of repair template used is as follows: 1.6 µM for *spo-11(ΔC21)::3xFLAG*, 2.2 µM for *spo-11(ΔC39)::3xFLAG*. For *dsb-1(R116A)*, *3xFLAG::dsb-1(R116A)*, *dsb-1(R116A) spo-11(ΔC21)*, *dsb-1(RD)*, *3xFLAG::dsb-1(RD), dsb-1(RD) spo-11(3A)*, *3xFLAG::dsb-1 spo-11(ΔC39),* and *3xFLAG::dsb-1* strain generation, a 5 µL mixture containing 17.5 µM Cas9 protein, 17.5 µM trans-activating tracrRNA, 17.5 µM crRNA targeting *dpy-10* (1.4 µM) and *dsb-1* (16.1 µM), either 6 µM Ultramer (single-strand DNA repair template) or melted double-strand DNAs generated by PCR as a repair template and 0.5 µM repair template of *dpy-10* was injected into gonads of 24 hours post-L4 stage N2, *3xFLAG::dsb-1*, *spo-11(ΔC21)* or *spo-11(3A)* mutants. The concentration of repair template used is as follows: 0.08 µM for *dsb-1(R116A)*, *3xFLAG::dsb-1(R116A)* and *dsb-1(R116A) spo-11(ΔC21)*, 6 µM for *dsb-1(RD)*, *3xFLAG::dsb-1(RD)* and *dsb-1(RD) spo-11(3A)*, 0.5 µM for *3xFLAG::dsb-1 spo-11(ΔC39),* 0.2 µM for 3x*FLAG::dsb-1*. To prevent re-editing by the CRISPR Cas9, silent mutations were introduced into the DNA repair templates. Dpy or Rol F1 generation worms were individually picked to new plates to produce progenies and screened by PCR and DNA sequencing for successful editing. The list of crRNAs, genotyping primers, and synthesized repair templates are shown in **Supplemental Table S4**. Ultramers, crRNA and tracrRNAs were purchased from IDT (IDT, Coralville, LA), and Cas9 proteins were purchased either from IDT or University of California Berkeley QB3 MacroLab (Berkeley, CA).

### Western blot

For the immunoblots shown in **Supplemental Figure S4B, S5B, S5C, S6C** and **S6D**, 24 h post L4 adult worms were mixed with 4x SDS sample buffer and snap-frozen in liquid nitrogen. The samples were boiled at 95°C for 5 min and vortexed for 20 min. SDS-PAGE was performed using 7.5% SDS-PAGE gel (Fujifilm Wako 198-14941, Osaka, Japan), and proteins were transferred to a PVDF membrane (Millipore IPVH00010, Burlington, MA) at 4°C, 80 V for 2 h. The membrane was blocked with 2.5% skim milk (Nacalai Inc 31149-75, Kyoto, Japan) in TBST buffer at room temperature for 1h, and probed with primary FLAG antibody in TBST containing 2.5% skim milk at 4°C overnight. After washing three times for 10 min each with TBST, the membrane was incubated with secondary antibody (anti-mouse-HRP) in TBST containing 2.5% skim milk, followed by three additional washes for 10 min each. Chemi-Lumi one super (Nacalai Inc 02230-30, Kyoto, Japan) was used to visualize protein bands using an ImageQuant LAS4000 (Cytiva Marlborough, MA). The list of antibodies is shown in **Supplemental Table S3**. Band intensities were quantified using Fiji. For FLAG immunoblots, the relative protein abundance in each mutant was calculated by the following steps: first, signal intensities of corresponding regions in the untagged wild-type lane on the same membrane were subtracted from each FLAG signal to correct for background. Then, local background-corrected actin signal measurements were used to normalize the FLAG signals. Finally, the resulting mutant lane intensities were further normalized to the intensity of the corresponding tagged control band.

## Author Contributions

Conceptualization, A.S.-C., and P.M.C.; methodology, K.K., C.M.R.-R., A.S.-C., and P.M.C.; software, C.M.R.-R. and P.M.C.; investigation, K.K., L.Z., C.M.R.-R., L.M., X. L.,T.Y., A.S.-C., and P.M.C.; writing, K.K., L.Z., C.M.R.-R., L.M., X. L., A.S.-C., and P.M.C..; funding acquisition, X. L., A.S.-C. and P.M.C.

## Acknowledgments

We thank Corentin Claeys-Bouuaert, Bernard de Massy, Abby Dernburg, Scott Keeney, Lewis Stevens, Anne Villeneuve, and Judith Yanowitz for kindly discussing this research with us. Some strains were obtained from the Caenorhabditis Genetics Center, which is funded by the NIH Office of Research Infrastructure Programs (P40 OD010440) and the National Bioresource Project Japan (Shohei Mitani lab). We thank Adam Guy, Tomoko Nishiyama, Masahiro Takado, Chieko Wada, and Shige H. Yoshimura for providing strains, reagents or equipment used in this study. This work is supported by JSPS Kakenhi 25K09508, 22K06079 (A. S-C.), 26K23144, 24K01955 (P.M.C.), and 25KJ0201 (X.L.) and JST CREST JPMJCR24B4 (P.M.C).

---

**Figure S1.**
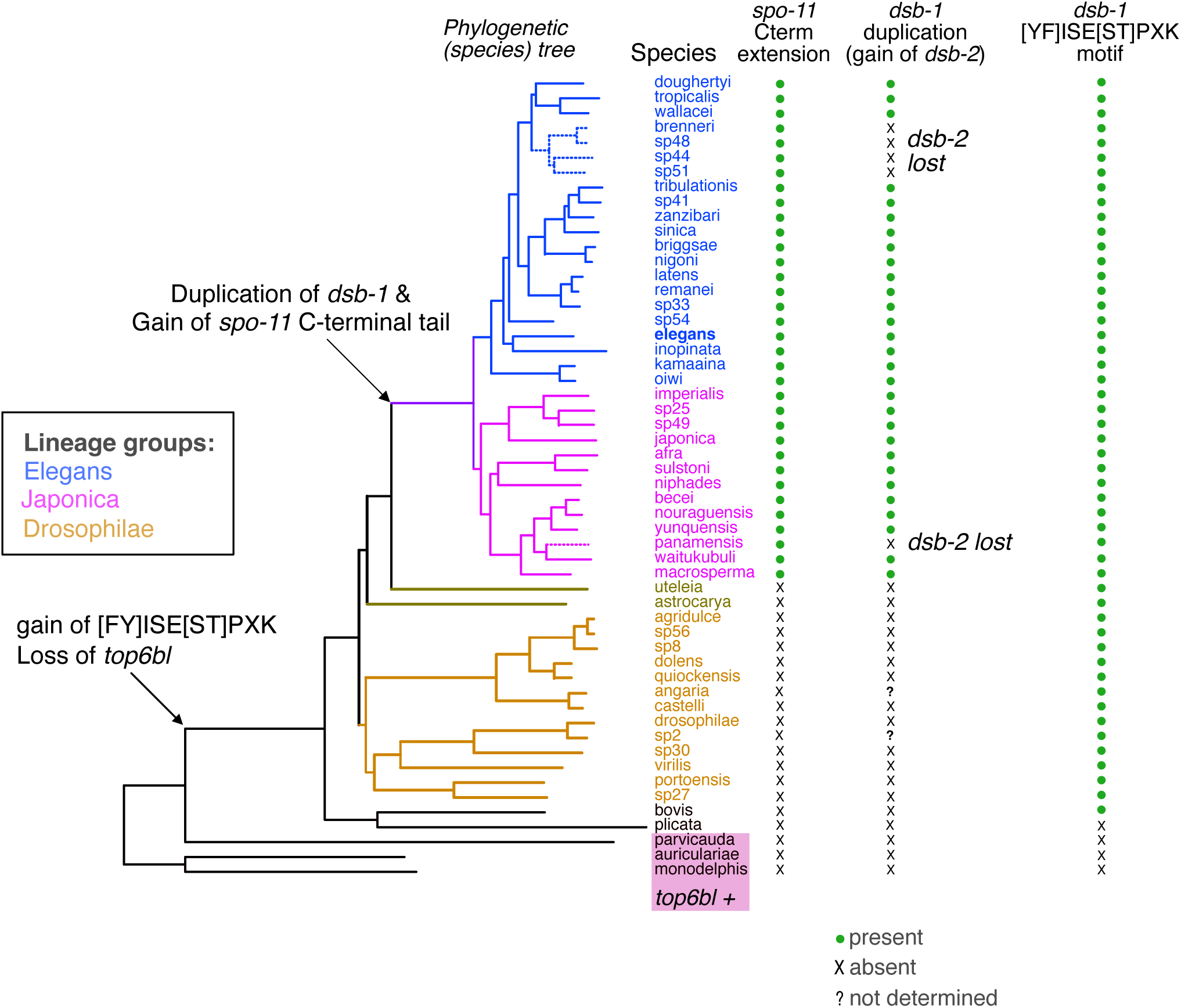
Mapping of *top6bl* presence, [YF]ISE[ST]PXK motif emergence, SPO-11C-terminal extension, and duplication of ancient Rec114 (*dsb-1*) onto the *Caenorhabditis* phylogeny (Left) Proteome-derived species phylogenetic tree of *Caenorhabditis* worms, with branch lengths proportional to the estimated number of amino acid substitutions per site. The three species retaining *top6BL* are highlighted in lavender. (Right) List of species shown with presence or absence of the indicated protein characteristics.

**Figure S2:**
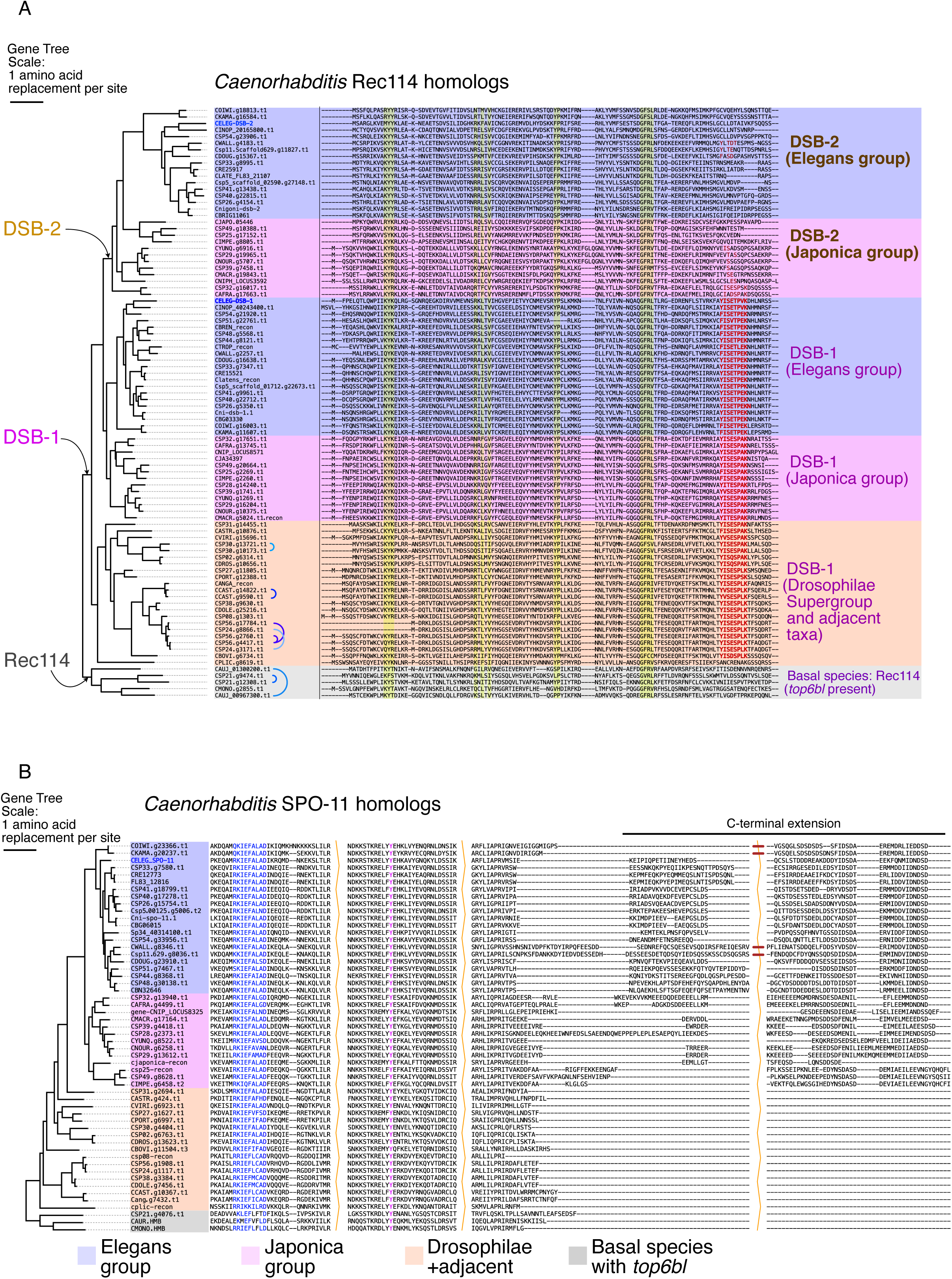
The SPO-11C-terminal extension correlates with an ancient duplication of *Rec114(dsb-1)* in the ancestor of the Elegans and Japonica groups (A) (Left) Gene tree of all identified *Caenorhabditis Rec114(dsb-1/2)* genes created with IQTree3 with branch lengths showing substitutions. (Right) Rec114(DSB-1/2) protein sequences of corresponding species. The [YF]ISE[ST]PXL motif is shown in red, four simple structural motifs shared throughout the family are highlighted in yellow, and *C. elegans* DSB-1 and DSB-2 labels are highlighted in blue. Species outside the Elegans and Japonica groups do not possess DSB-2, but some of them have recent duplications of DSB-1 (denoted by half circles next to the protein names). (B) (Left) Gene tree of all identified *Caenorhabditis spo-11* genes created with IQTree3, as in (A). (Right) Sequences from informative MSA regions of SPO-11 with elisions indicated by orange zigzags. From left, the regions show the N-terminal RKIEFALAD motif predicted to be involved in DSB-1 binding (coordinates around 50aa in *C. elegans*, shown in blue); the catalytic tyrosine residues (at 119aa in *C. elegans,* denoted in shown), then the C-terminal extension with two elisions involving four species (indicated by dark bars) with longer sequences, omitted here for presentation. The label "recon" in all protein names indicates manual reconstruction from the genome (see Methods).

**Figure S3:**
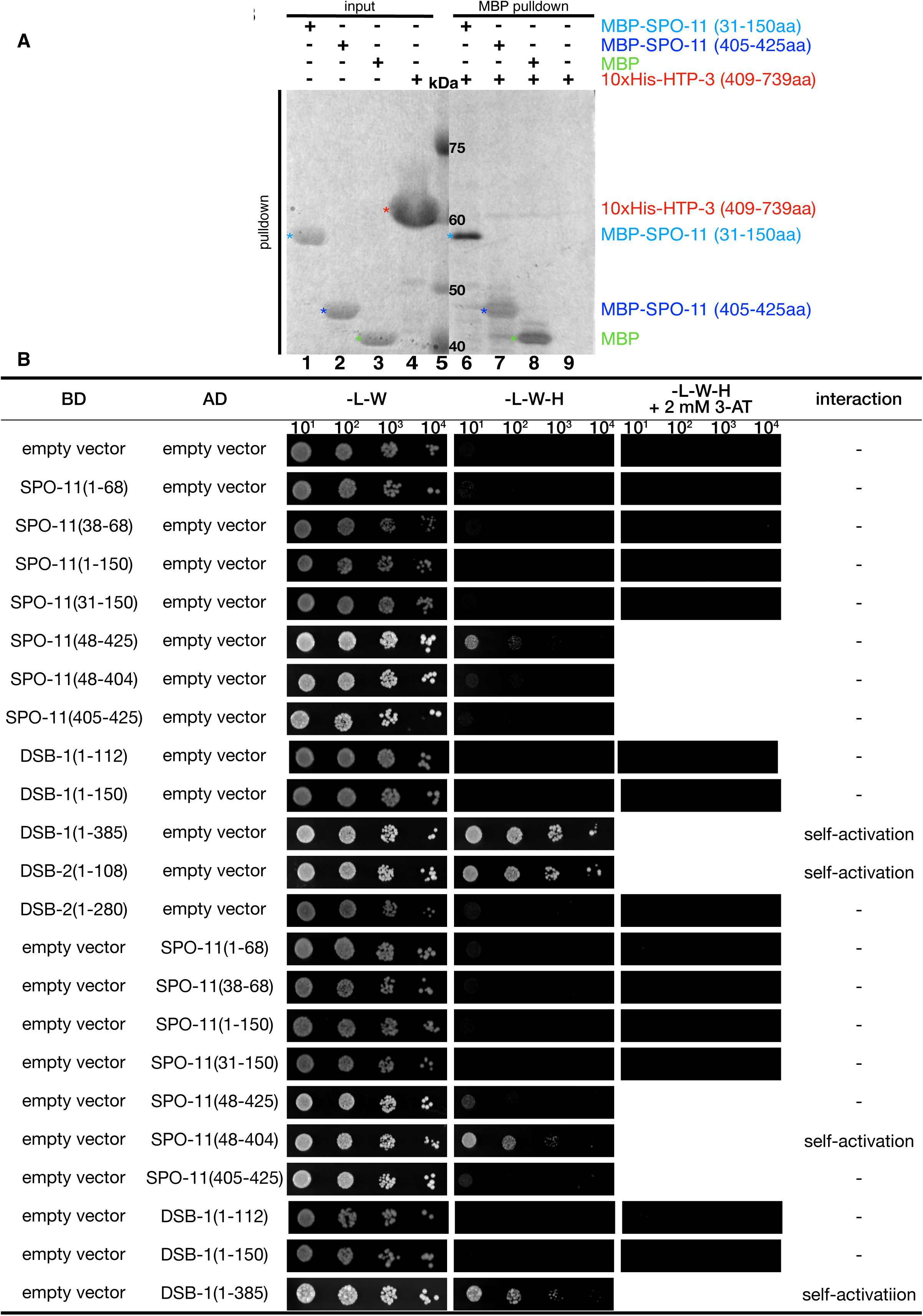
Controls for pulldown and Y2H assays. (A) The negative control 10xHis-tagged HTP-3 (409-739aa fragment) was not pulled down by MBP-tagged SPO-11 N-terminal (lane 6), C-terminal (lane 7) fragments nor MBP (lane 8), indicating that 10xHis-tag itself does not interact with MBP or MBP fused SPO-11 fragments. Proteins in the input (lanes 1-5) were visualized by Coomassie Brilliant Blue staining and proteins in the eluates (lanes 6-9) were visualized by silver-staining. **(B)** All Y2H results of indicated protein fragments with 10-fold to 10^4^-fold serial dilutions. Colony growth was assessed on medium lacking leucine (L), tryptophan(W) and/or histidine (H), and with 2 mM 3-amino-1,2,4-trizole (3-AT).

**Figure S4.**
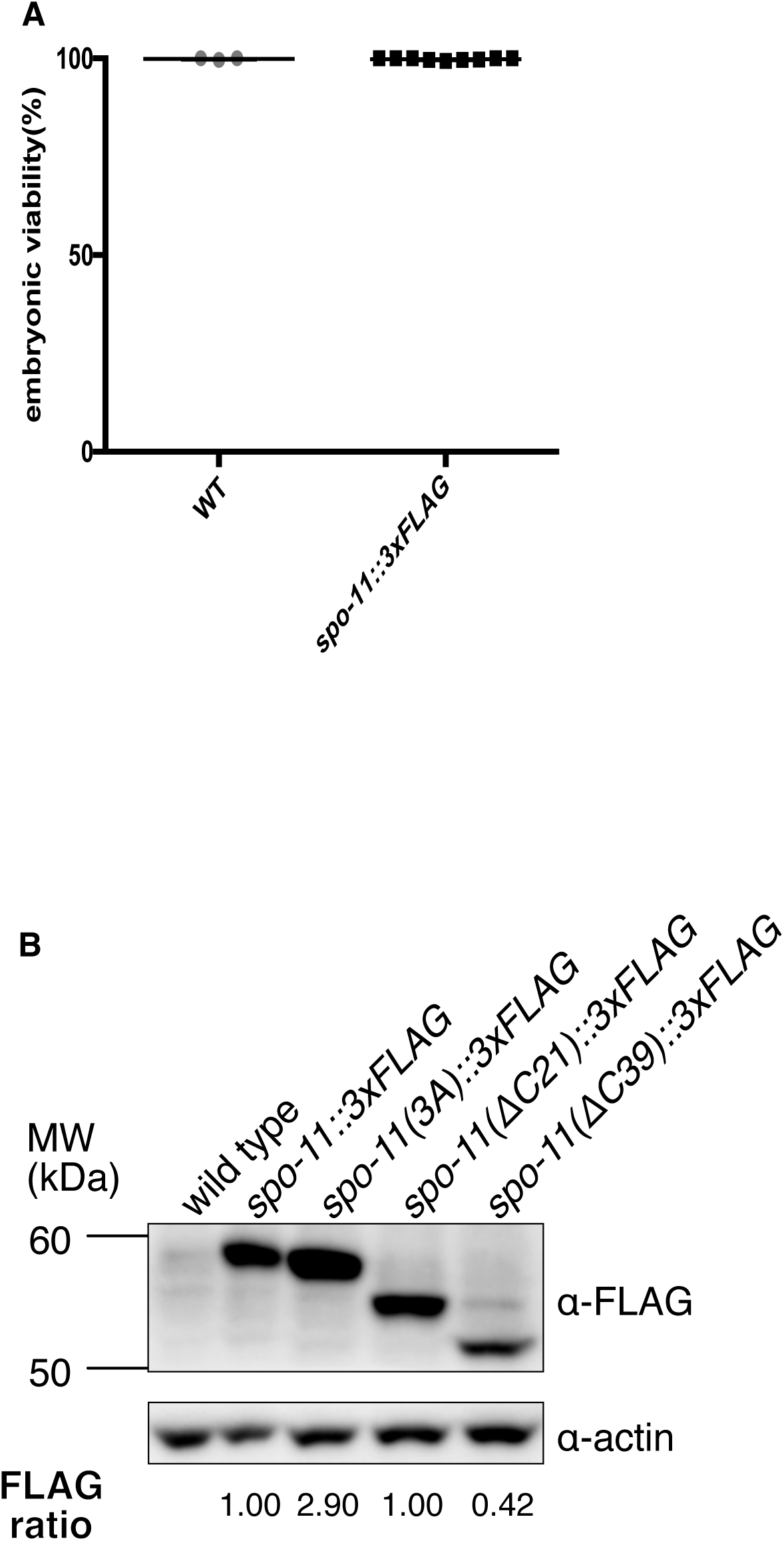
Embryonic viability analysis, assessment of protein levels by Western blot and DSB level quantification by RAD-51 immunofluorescence of *spo-11* mutants as well as FLAG-tagged control strains. **(A)** FLAG tagging of SPO-11 did not interfere with embryonic viability. Embryonic viability of N2 wild type and *spo-11:3x:FLAG(wt)*. Data are presented as percentages. Statistical significance was assessed using a two-tailed t-test with Welch’s correction. **(B)** Western blot of FLAG-tagged SPO-11 probed with an anti-FLAG antibody. FLAG-tagged SPO-11 proteins were detected in all FLAG-tagged strains (100 worms, 24hr post-L4 stage). Anti-actin immunoblotting is shown as a loading control. Values shown below the bands represent FLAG signals that have been background-corrected using the untagged wild-type control, normalized to actin, and normalized to the corresponding FLAG-tagged wild-type control (n=2).

**Figure S5:**
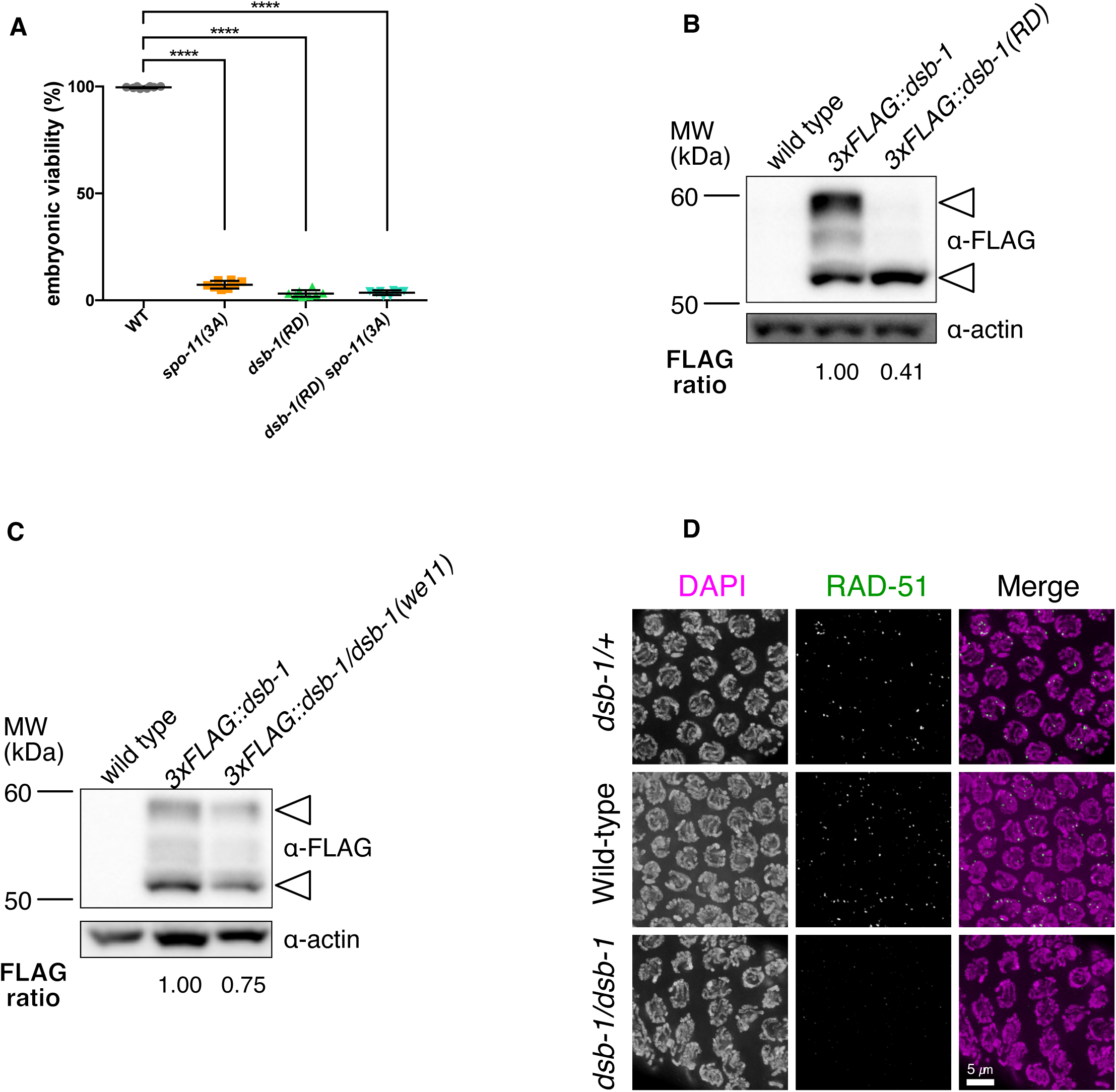
Assessment of the contribution of reduced DSB-1 protein abundance to mutant phenotypes (A) Embryonic viability of *spo-11(3A)*, *dsb-1(RD)* and *dsb-1(RD) spo-11(3A)*. Data are presented as percentages. Statistical significance was assessed using a two-tailed t-test with Welch’s correction, ****p<0.0001. **(B)** Western blot of FLAG-tagged DSB-1 probed with an anti-FLAG antibody. FLAG-tagged DSB-1 proteins were detected in *3xFLAG::dsb-1(wt)* and *3xFLAG::dsb-1(RD)*(100 worms, 24 hr post-L4 stage). Arrowheads indicate the two specific bands detected in the blot. Anti-actin immunoblotting is shown as a loading control. Values shown below the bands represent FLAG signals background-corrected using the untagged wild type control, normalized to actin, and normalized to the corresponding FLAG-tagged wild-type control (n=2). **(C)** Western blot of FLAG-tagged DSB-1 probed with an anti-FLAG antibody. FLAG-tagged DSB-1 proteins were detected in *3xFLAG::dsb-1(wt)* and *3xFLAG::dsb-1(wt)/dsb-1(we11)* heterozygous(100 worms, 24 hr post-L4 stage). Arrowheads indicate the two specific bands detected in the blot. Anti-actin immunoblotting is shown as a loading control. Values shown below the bands represent FLAG signals background-corrected using the untagged wild type control, normalized to actin, and normalized to the corresponding FLAG-tagged wild type control(n=3). **(D)** Representative immunofluorescence images of RAD-51 foci in mid-pachytene oocytes (zone 4) of the gonads for each genotype indicated. Scale bar, 5 µm.

**Figure S6:**
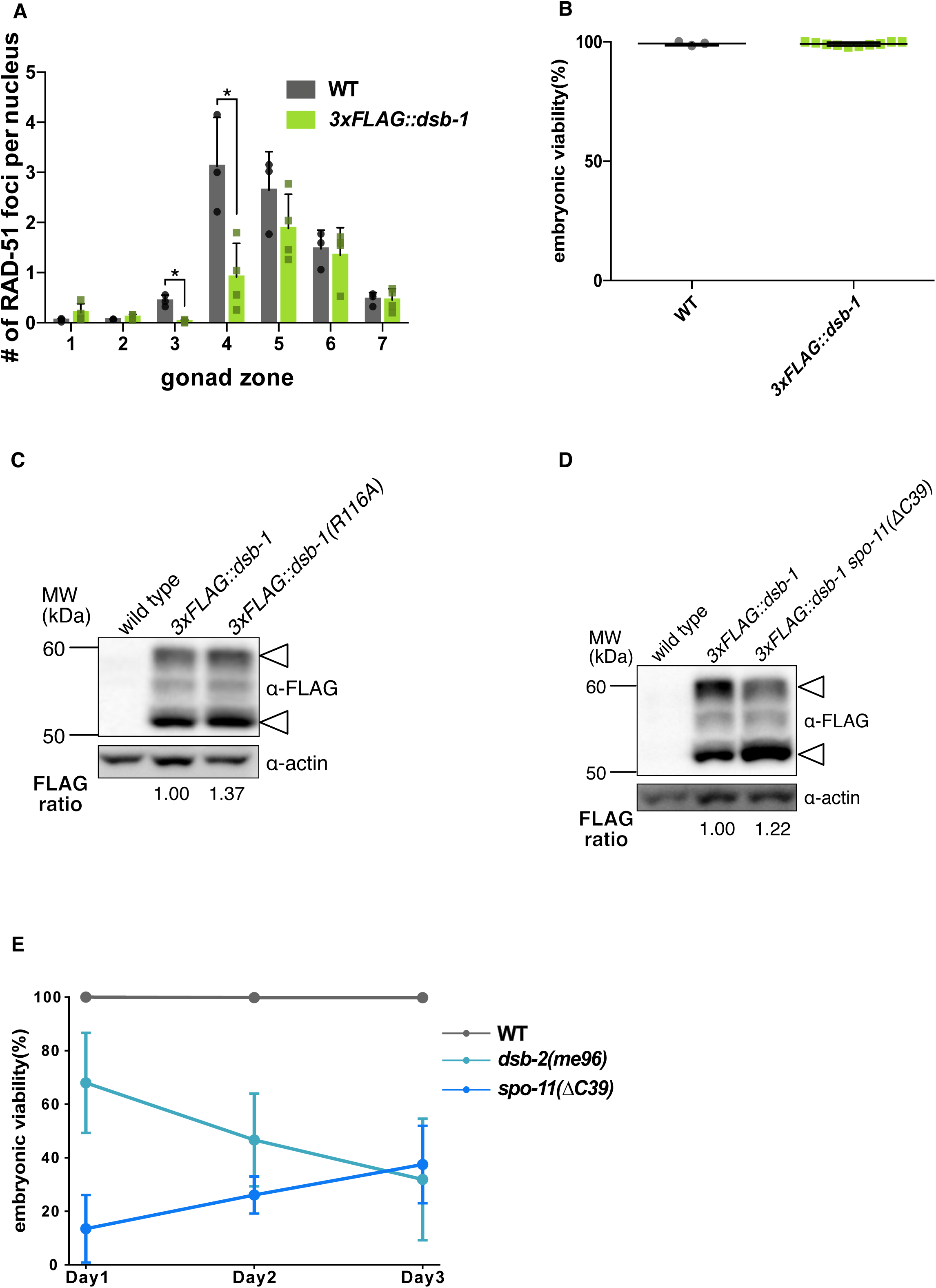
Embryonic viability analysis, assessment of protein levels by Western blot and DSB level quantification by RAD-51 immunofluorescence of *dsb-1* mutants as well as FLAG-tagged control strains. **(A)** Quantification of RAD-51 foci in the gonads in WT and *3xFLAG::dsb-1(wt)*. Data are presented as mean±SD (bar, overall mean; dot, mean of each individual gonad; The numbers of scored nuclei in zone 1-7 were as follows: for wild type (3 gonads), 126, 141, 135, 124, 120, 99, 80; for *3xFLAG::dsb-1(wt)* (4 gonads), 130, 139, 173, 192, 207, 169, 135. Statistical significance was assessed using a two-tailed t-test with Welch’s correction, *p<0.05. **(B)** Embryonic viability of *3x:FLAG::dsb-1(wt)*. Data are presented as percentages. Statistical significance was assessed using a two-tailed t-test with Welch’s correction. **(C)** Western blot of FLAG-tagged DSB-1 probed with an anti-FLAG antibody. FLAG-tagged DSB-1 proteins were detected in *3xFLAG::dsb-1(wt)* and *3xFLAG::dsb-1(R116A)* (100 worms, 24hr post-L4 stage). Arrowheads indicate the two specific bands detected in the blots. Anti-actin immunoblotting is shown as a loading control. Values shown below the bands represent FLAG signals background-corrected using the untagged wild-type control, normalized to actin, and normalized to the corresponding FLAG-tagged wild-type control (n=3). **(D)** Western blot of FLAG-tagged DSB-1 probed with an anti-FLAG antibody. FLAG-tagged DSB-1 proteins were detected in *3xFLAG::dsb-1(wt)* and *3xFLAG::dsb-1(wt) spo-11(ΔC39)* (100 worms, 24hr post-L4 stage). Arrowheads indicate the two specific bands detected in the blot. Anti-actin immunoblotting is shown as a loading control. Values shown below the bands represent FLAG signal quantitation: background-corrected using the untagged wild-type control, normalized to actin, and normalized to the corresponding FLAG-tagged wild-type control (n=1). **(E)** Embryonic viability of self-fertilized progeny from hermaphrodites of the indicated genotypes during the indicated time intervals after the L4 larval stage. Data are presented as mean ± SD (WT: n = 5; *dsb-2(me96)*: n = 9; *spo-11(ΔC39)*: n = 10).

